# Comparative mapping of single-cell transcriptomic landscapes in neurodegenerative diseases

**DOI:** 10.1101/2024.12.13.628436

**Authors:** E. Keats Shwab, Zhaohui Man, Daniel C. Gingerich, Julia Gamache, Melanie E. Garrett, Geidy E. Serrano, Thomas G. Beach, Gregory E. Crawford, Allison E. Ashley-Koch, Ornit Chiba-Falek

**Author notes:** To whom correspondence should be addressed: Ornit Chiba-Falek Division of Translational Brain Sciences, Dept of Neurology Duke University School of Medicine Durham, North Carolina 27710, USA. These authors contributed equally to this study (E. Keats Shwab, Zhaohui Man). **LIST OF ABBREVIATIONS:** AD (Alzheimer’s disease), PD (Parkinson’s disease), DLB (dementia with Lewy bodies), NDD (neurodegenerative disease), NFT (neurofibrillary tangle), fPD (familial PD), GWAS (genome-wide association study), snRNA-seq (single-nucleus RNA sequencing), TC (temporal cortex), NC (normal control), QC (quality control), OPC (oligodendrocyte precursor cell), DEG (differentially expressed gene), PMI (postmortem interval), FDR (false discovery rate), TF (transcription factor), Astro (astrocyte), Exc (excitatory neuron), Inh (inhibitory neuron), Micro (microglia), Oligo (oligodendrocyte), PCA (principal component analysis), UMAP (uniform manifold approximation and projection), ER (endoplasmic reticulum), APP (amyloid precursor protein), SN (substantia nigra), KPBBB (Kathleen Price Bryan Brain Bank), BSHRI (Banner Sun Health Research Institute), USSLB (Unified Staging System for Lewy Body Disorders), IRB (institutional review board), NIH (National Institutes of Health), NINDS (National Institute of Neurological Disorders & Stroke), NIA (National Institute on Aging).

## Abstract

**INTRODUCTION:** Alzheimer’s disease (AD), Dementia with Lewy bodies (DLB), and Parkinson’s disease (PD) represent a spectrum of neurodegenerative disorders (NDDs). Here, we performed the first direct comparison of their transcriptomic landscapes.

**METHODS:** We profiled the whole transcriptomes of NDD cortical tissue by snRNA-seq. We used computational analyses to identify common and distinct differentially expressed genes (DEGs), biological pathways, vulnerable and disease-driver cell subtypes, and alteration in cell-to-cell interactions.

**RESULTS:** The same vulnerable inhibitory neuron subtype was depleted in both AD and DLB. Potentially disease-driving neuronal cell subtypes were present in both PD and DLB. Cell-cell communication was predicted to be increased in AD but decreased in DLB and PD. DEGs were most commonly shared across NDDs within inhibitory neuron subtypes. Overall, we observed the greatest transcriptomic divergence between AD and PD, while DLB exhibited an intermediate transcriptomic signature.

**DISCUSSION:** These results help explain the clinicopathological spectrum of this group of NDDs and provide unique insights into the shared and distinct molecular mechanisms underlying the pathogenesis of NDDs.

## 1. BACKGROUND

Age-associated neurodegenerative diseases (NDD) such as Alzheimer’s disease (AD), Parkinson’s disease (PD), and Dementia with Lewy bodies (DLB) exhibit overlapping molecular pathologies (Fig. 1A). For example, Lewy bodies are present in more than half of AD cases^1,2^, and tau neurofibrillary tangles (NFTs) have been identified in brains of patients with familial PD (fPD)^3^. Tau also appears to be a common component of Lewy bodies in association with SNCA^4,5^. Tau NFTs and Aβ plaques are also associated with DLB in approximately 70% of cases^6,7^, indicating convergence of underlying pathological mechanisms of both AD and PD in DLB^8^. Evidence suggests that these co-pathologies of tau, Aβ and SNCA aggregates are not merely coincidental but that these molecules are also likely involved in seeding the aggregation of one another^9^.

**Figure 1.**
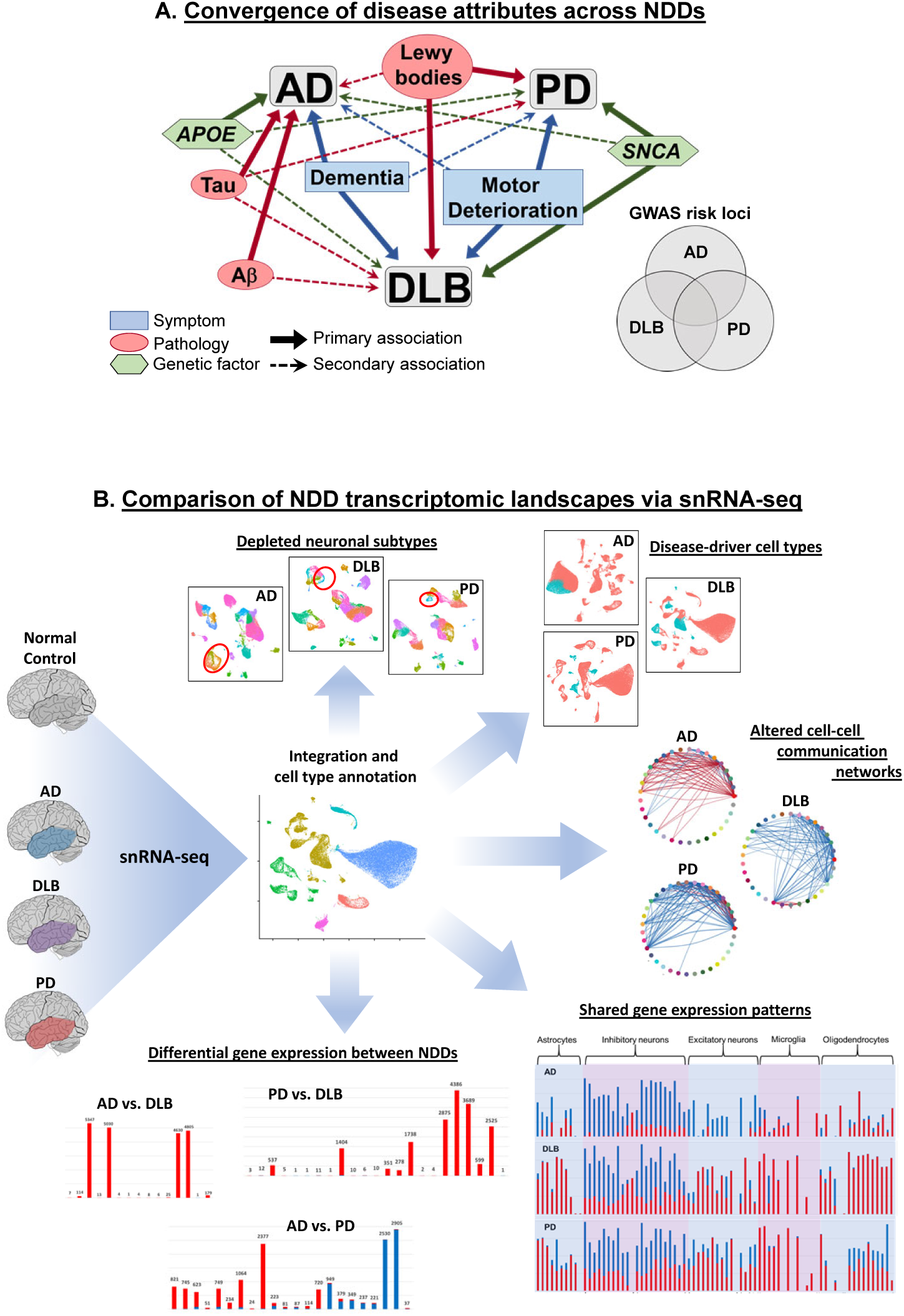
Elucidating similarities and differences in transcriptomic landscapes underlying shared and distinct pathologic attributes of AD, PD, and DLB. A. Convergence of disease attributes across NDDs. Dementia is a defining symptom of both AD and DLB but may also be present in PD, while motor deterioration is a primary symptom of PD and DLB but may also be present in AD. Lewy bodies are a hallmark of both PD and DLB, but are also present in over half of AD cases, while tau and Ab, hallmarks of AD, are often present in DLB, and tau is a common component of Lewy bodies. *APOE* variants represent the highest genetic risk factor for AD, but mutations have also been linked to DLB risk and cognitive decline in PD. *SNCA* is primarily associated with PD and DLB, but mutations in this gene are also are associated with increased risk of AD. Furthermore, numerous GWAS identified risk alleles show overlap across all three NDDs. B. Comparison of NDD transcriptomic landscapes via snRNA-seq. TC samples from 12 donors diagnosed with AD, DLB, and PD, as well as normal controls, were used for snRNA-seq analysis, followed by integration of transcriptomic datasets and cell type annotation. Datasets were examined for depletion of neuronal cell subtypes in each NDD compared to NC nuclei, identification of disease-driver cell types with enriched expression of GWAS genes, changes in cell-to-cell communication between cell subtypes in NDD and NC nuclei, shared genes differentially expressed in each NDD compared to NC nuclei, and differential gene expression between each pair of NDDs.

In addition to co-pathologies, commonalities are evident in the underlying genetic architectures of these three NDDs. Genome-wide association studies (GWAS) focusing on each of these NDDs have identified variants separately associated with increased risk for AD^10–12^, PD^13,14^, and DLB^15,16^, and overlap in genetic risk factors between the NDDs has also been observed. For example, mutations in *APOE*, the primary risk factor for AD, have also been linked to increased risk of DLB^15^ and cognitive decline in PD^17^. Additionally, *SNCA* mutations have been similarly linked to both AD^18^ and DLB^15^ risk. Furthermore, mutations in GWAS AD risk genes including *APP*^19^, *PSEN1*^19–21^, and *PSEN2*^19,21^, and GWAS PD risk genes including *LRRK2*^22^, *MAPT*^23^, and *SCARB2*^24^ have also been experimentally linked to DLB. However, numerous loci with positive risk correlations for either AD or PD are not correlated with DLB risk^15^. These data indicate unique as well as shared genetic underpinnings for each of these NDDs.

The majority of disease-associated SNVs are located in noncoding genomic regions, suggesting that in many cases allelic disease effects may derive from altered gene regulation rather than altered protein coding, and therefore additional information is required to determine the relevant target genes impacted. The development of single-cell transcriptomics has helped elucidate these changes in gene regulation by enabling examination of expression patterns within individual brain cells of NDD patients at an unprecedented cell type and subtype resolution. This methodology has been applied individually to AD^25–28^ and PD^29,30^, but for DLB only bulk transcriptomic studies have been previously performed^31–33^. Moreover, no study to date has compared the transcriptional profiles across these NDDs. In this work, we used single-nucleus RNA sequencing (snRNA-seq) to directly compare and contrast for the first time the transcriptomic signatures of these three prevalent NDDs in order to elucidate shared and unique dysregulated genes and networks among these pathologies (Fig. 1B). We compared gene expression in 12 temporal cortex (TC) samples of donors diagnosed with each NDD: AD, PD and DLB, to 12 neurologically normal control (NC) samples. Moreover, we performed examinations of differential gene expression between each pair of NDDs (i.e. AD vs. PD, AD vs. DLB, PD vs. DLB), all at a granular cell subtype level of precision. We furthermore identified and characterized specific cell subtypes depleted in each of the NDDs, and predicted changes in cell-to-cell communication patterns associated with each disorder. Our findings yield novel insights into pathology-associated changes in gene expression that may facilitate the development of new detection and treatment strategies targeting specific NDDs or potentially effective in the treatment of a range of disorders.

## 2. METHODS

### 2.1 Human post-mortem brain tissue samples

The demographics, pathological notes, and other metadata for this study cohort are detailed in Table S1. Extensive pathology information for PD samples is provided in Table S2. Frozen human TC tissue samples from donors clinically diagnosed with AD (*n* = 12), DLB (*n* = 12) and NC donor samples (*n* = 12) were obtained from the Kathleen Price Bryan Brain Bank (KPBBB) at Duke University. Samples from donors diagnosed with PD (*n* = 12) were obtained from the Banner Sun Health Research Institute (BSHRI)^88^. Normal controls were derived from donors with no clinical history of neurological disorder and samples had no neuropathological evidence of neurodegenerative diseases. Clinical diagnosis of AD was pathologically confirmed using Braak staging (AT8 immunostaining) and amyloid deposition assessment (4G8 immunostaining) for all AD samples. All AD tissue donors were in Braak & Braak Stage III-V. DLB clinical diagnoses were pathologically confirmed based on criteria described by McKeith et al.^7^ All DLB donors were confirmed to exhibit Lewy-related pathology within the neocortical, limbic, or brainstem regions and showed low levels of AD neuropathologic change (Braak stages I or II), with the exception of donor 1097 which exhibited Braak stage III pathology. Donor patient PD diagnoses were defined by the presence of two of the three cardinal clinical signs of resting tremor, muscular rigidity and bradykinesia. Additionally, diagnoses of all PD samples were confirmed in autopsy by observation of pigmented neuron loss and the presence of Lewy bodies in the SN. Neuropathological states of PD samples were confirmed *postmortem* using established clinical practice recommendations for McKeith scoring^83^ and staging via the Unified Staging System for Lewy Body Disorders (USSLB)^89^. All PD samples for which information was available had McKeith scores ranging from moderate to severe (2-4) in both the amygdala and SN. Where available, TC McKeith scores for most of the PD samples were either 0-1, with one sample each receiving scores of 2 and 3, indicating mild or absent PD pathology in this region for the majority of samples. USSLB stages of PD samples ranged from II-IV. PD samples 96-36 and 96-49 were lacking specific USSLB stage determination due to harvesting prior to BSHRI standardization of stage determination protocol. All tissue donors were Caucasians with the *APOE* e3/e3 genotype. The project was approved for exemption by the Duke University Health System Institutional Review Board. The methods described were conducted in accordance with the relevant guidelines and regulations.

### 2.2 Nuclei isolation from post-mortem human brain tissue

The nuclei isolation procedure has been described^28^, and was based on previous studies^90,91^ and optimized for single-cell experiments. 100-200 mg of human TC brain tissue samples were thawed in Lysis Buffer (0.32 M Sucrose, 5 mM CaCl_2_, 3 mM Magnesium Acetate, 0.1 mM EDTA, 10 mM Tris-HCl pH 8, 1 mM DTT, 0.1% Triton X-100) and homogenized with a 7 ml dounce tissue homogenizer (Corning) and filtered through a 100 μm cell strainer, transferred to a 14 x 89 mm polypropylene ultracentrifuge tube, and underlain with sucrose solution (1.8 M Sucrose, 3 mM Magnesium Acetate, 1 mM DTT, 10 mM Tris-HCl, pH 8). Nuclei were separated by ultracentrifugation for 15 minutes at 4°C at 107,000 RCF. Supernatant was aspirated, and nuclei were washed with 1 ml Nuclei Wash Buffer (10 mM Tris-HCl pH 8, 10 mM NaCl, 3 mM MgCl_2_, 0.1% Tween-20, 1% BSA, 0.2 U/μL RNase Inhibitor). Resuspended nuclei were centrifuged at 300 RCF for 5 minutes at 4°C, and supernatant was aspirated. Nuclei were then resuspended in Wash and Resuspension Buffer (1X PBS, 1% BSA, 0.2 U/μL RNase Inhibitor), then filtered through a 35 μm strainer. Nuclei concentrations were determined using a Countess™ II Automated Cell Counter (ThermoFisher) and nuclei quality was assessed at 10X and 40X magnification using an Evos XL Core Cell Imager (ThermoFisher).

### 2.3 snRNA-seq library preparation and sequencing

snRNA-seq libraries were constructed as previously^28^ using the Chromium Next GEM Single Cell 3’ GEM, Library, and Gel Bead v3.1 kit, Chip G Single Cell kit, and i7 Multiplex kit (10X Genomics) according to manufacturer’s instructions. For each sample, 10,000 nuclei were targeted. Library quality control was performed on a Bioanalyzer (Agilent) with the High Sensitivity DNA Kit (Agilent) according to manufacturer’s instructions and the 10X Genomics protocols. Libraries were submitted to the Sequencing and Genomic Technologies Shared Resource at Duke University for quantification using the KAPA Library Quantification Kit for Illumina® Platforms and sequencing. Groups of four snRNA-seq libraries were pooled on a NovaSeq 6000 S1 50bp PE full flow cell to target a sequencing depth of 400 million reads per sample (Read 1 = 28, i7 index = 8, and Read 2 = 91 cycles). Sequencing was performed blinded to age, sex, and diagnosis.

### 2.4 snRNA-seq data processing

Raw snRNA-seq sequencing data were converted to FastQ format, aligned to a GRCh38 pre-mRNA reference, filtered, and counted using CellRanger 4.0.0 (10X Genomics). Subsequent processing was done using Seurat 4.0.1^92^. Filtered feature-barcode matrices were used to generate Seurat objects for the individual samples. For QC filtering, nuclei below the 1^st^ and above the 99^th^ percentile for number of features were excluded. Nuclei above the 95^th^ percentile for mitochondrial gene transcript proportion (or >5% mitochondrial transcripts if 95^th^ percentile mitochondrial transcript proportion was <5%) were also excluded. Because experiments were conducted in nuclei rather than whole cells, mitochondrial genes were subsequently removed. The individual sample Seurat objects were merged into one, and were iteratively normalized using SCTransform^93^ with glmGamPoi, which alleviates bias from weakly-expressed genes^94^. Batch correction was performed using reference-based integration^34^ on the individual sample normalized datasets, which improves computational efficiency for integration.

### 2.5 Doublet/Multiplet detection in snRNA-seq data

Multiplets comprising different cell types (heterotypic) were excluded from snRNA-seq data by considering the “hybrid score”, as described previously^28^. The hybrid score is calculated as (x_1_ – x_2_) / x_1_, where x_1_ is the highest and x_2_ is the second highest prediction score^95^. Heterotypic multiplets would be expected to exhibit competing cell type prediction scores due to the presence of transcriptomic/epigenomic profiles from multiple cell types. Multiplets composed of one cell type (homotypic) were identified based on the number of features per cell. snRNA nuclei with feature counts > 99^th^ percentile were excluded. Removal of homotypic multiplets in this manner is expected to also aid in filtering of heterotypic multiplets.

### 2.6 Cell type and subtype cluster annotation

Cell type annotation was conducted using a label transfer method^34^ and a previously annotated reference dataset from human M1. Batch-corrected data from both our dataset and the human M1 dataset were used for label transfer. Nuclei with maximum prediction scores of <0.5 were excluded. Nuclei with a percent difference of <20% between first and second highest cell type prediction scores were termed “hybrid” and excluded^95^. Endothelial cells and VLMCs were in low abundance and did not form distinct UMAP clusters and were thus excluded. Following PCA, dimensionality was examined using an Elbow plot and by calculating variance contribution of each PC. UMAP was then run using the first 30 PCs, and nuclei were clustered based on UMAP reduction. The resolution levels for cluster delineation were selected after comparison of a range of values as it was determined to provide optimal distinction between populations of nuclei displaying unique gene expression profiles as evidenced by their separation from one another in UMAP space. Counts of predicted major cell types based on the label transfer were examined for each of the clusters, and clusters were manually annotated based on the majority cell type for each cluster (e.g., ‘Exc1’, ‘Exc2’, etc.).

### 2.7 Human M1 reference data processing

To optimize label transfer, we re-processed previously published human primary motor cortex (M1) snRNA-seq data^96^ to map it to GRCh38 Ensembl 80 as we did with our data^28^. FastQ files were obtained from the Neuroscience Multi-omic Data Archive (NeMO: https://nemoarchive.org/) and were aligned to the same GRCh38 pre-mRNA reference used for our data, filtered, and counted using CellRanger 4.0.0 (10X Genomics). Filtered feature-barcode matrices were used to generate separate Seurat objects for each sample, with nuclei absent from the annotated metadata excluded. Seurat objects were merged and iteratively normalized using SCTransform^93^ with glmGamPoi. Batch correction was performed using reference-based integration^34^ on the normalized datasets. The 127 transcriptomic cell types in this data were grouped into 8 broad cell types including astrocytes, endothelial cells, excitatory neurons, inhibitory neurons, microglia, oligodendrocytes, OPCs, and VLMCs.

### 2.8 Covariate selection for differential analyses

Prior to differential analysis, as previously described, ^28^ we estimated the impact of multiple technical variables as well as donor-level characteristics separately for the snRNA-seq experiments (Table S1). Read counts were summed for all nuclei in each donor sample, resulting in only one expression value per sample per gene, as all nuclei from a particular donor would have identical donor characteristics. Genes with no expression for >20% of samples were subsequently removed, and all values were mean-centered and scaled prior to covariate analysis. PCA was then performed for genes using *prcomp* in R. We then carried out linear regression using *glm* in R for PCs explaining >10% of the variability in global expression on both nuclei- and donor-specific metadata variables to identify factors that should be included as covariates in differential analyses. Specifically, we selected the variable most associated (surpassing Bonferroni correction for multiple testing, *q*<0.05) with PC1 (or alternatively, the PC explaining the most variability) and regressed all genes on the associated variable to obtain gene residuals that are adjusted for its effect. We then performed PC analysis on the gene residuals, and in an iterative process, repeating the above steps until no additional metadata variables were associated with global expression (*q*<0.05). Following this process, age, sex, PMI, number of nuclei after QC filtering, median genes per cell, and average library size were selected as covariates for differential expression gene analysis.

### 2.9 Cell type proportion comparisons

To assess the selective loss of neuronal subtypes in each neurodegenerative disorder, we performed a depletion analysis using a beta regression model implemented in the glmmTMB package in R. The proportion of each neuronal subtype within each sample was calculated, and the association between the proportion and disease status was examined while adjusting for potential confounding variables such as age, sex, post-mortem interval (PMI), and the number of nuclei after filtering. The significance of the depletion was determined based on the Benjamini-Hochburg (FDR) adjusted p-values derived from the beta regression model.

### 2.10 Marker gene identification

To identify genes differentially expressed between depleted neuronal subtypes in each disease condition, we utilized the *FindMarkers* function from the Seurat package. The analysis was performed using a likelihood-ratio test, adjusting for latent variables including age, sex, PMI, and the number of nuclei after filtering. The gene expression comparison was made between the depleted neuronal subtypes in the disease samples and their corresponding subtypes in the control samples. Genes with a Benjamini-Hochburg (FDR) adjusted p-value less than 0.05 were considered significantly differentially expressed. The differentially expressed genes were further categorized into upregulated and downregulated genes based on their average log2 fold change.

### 2.11 Differential expression analysis

In order to identify DEGs at both the cell type and subtype levels between samples within our snRNA-seq dataset, we employed the NEBULA algorithm^45^. Specifically, the NEBULA-HL method was used as this process is optimized for estimating both nucleus-level and donor-level data overdispersions^45,97^. Prior to running NEBULA, for each cell type and cluster, genes expressed in less than 10% of cells in either group (PD or Normal) were filtered out. Age, sex, PMI, number of nuclei after QC filtering, median genes per cell, and average library size were included as fixed effects for NEBULA and sample donor ID was included as a random effect. Benjamini–Hochberg (FDR) correction for multiple testing was applied at the gene level to NEBULA-derived *p*-values. Adjusted *p*-values < 0.05 were deemed significant.

### 2.12 Vulnerable cell type identification

For each broad cell type in each disorder, DEGs were identified using the NEBULA algorithm as described above. GWAS-associated genes for each disorder were obtained from published studies, considering genes located within 500 kilobases upstream or downstream of the GWAS SNP chromosome locus. To create gene sets representing the convergence of genetic risk factors and cell type-specific dysregulation, we intersected the GWAS-associated genes with the DEGs identified for each broad cell type in each disorder. The resulting gene sets were considered as the putative driving forces or risk factors for the corresponding disorder. The vulnerability of each cell subtype to the disorder-specific gene sets was assessed using the AUCell package in R. For each cell subtype in each disorder, the following steps were performed:

1. The scRNA-seq data were subsetted to include only the cells belonging to the specific cell subtype.
2. The gene expression matrix was normalized and log-transformed.
3. The AUCell algorithm was applied to calculate the enrichment of the disorder-specific gene set in each cell, resulting in an AUC (Area Under the Curve) score for each cell.
4. Cells were assigned to a “vulnerable” or “non-vulnerable” group based on the AUC score threshold determined using the AUCell_exploreThresholds function.

To identify marker genes associated with the vulnerable cell subtypes, differential gene expression analysis was performed using the FindMarkers function in Seurat. The analysis was conducted between the vulnerable and non-vulnerable cells within each cell subtype, controlling for potential confounding variables. Genes with an FDR-adjusted *p*-value < 0.05 were considered significantly differentially expressed and were classified as marker genes.

### 2.13 Differential cell-to-cell communication

To investigate the role of cell-cell communication in the progression of neurodegenerative disorders (NDDs), we used CellChat, an R package for inference and analysis of intercellular communication networks from single-cell RNA sequencing (scRNA-seq) data^43^. CellChat integrates scRNA-seq data with a curated database of ligand-receptor interactions to quantify communication probabilities between cell populations and identify significant interactions. For each disease-normal pair, we created separate CellChat objects using the normalized data matrix and cell type annotations. We then applied CellChat functions to identify over-expressed genes and interactions, compute communication probabilities, and filter interactions. The inferred communication networks were stored in the CellChat object. To visualize the differences in cell-cell communication between disease and normal conditions, we employed CellChat’s plotting functions.

### 2.14 Biological pathway enrichment analysis

In order to understand the biological significance of gene sets derived from differential expression analyses, we employed the Metascape^41^ algorithm (https://www.metascape.org). The gene set of interest was input as the target gene list, and the total set of genes examined in the corresponding differential expression analysis was input as the background gene list. GO terms were considered significantly enriched with a fold-enrichment of at least 1.5 and an FDR-corrected enrichment *p*-value < 0.01. In order to group the enriched Metascape output GO terms into broader biological categories, Kappa similarities were determined for each pair of enriched GO terms, forming trees of hierarchical associations between terms, which were then used to delineate clusters of related terms. We then qualitatively assigned a major functional category label to each cluster based on assessment of common biological processes represented by the clustered GO terms.

### 2.15 Genome version and coordinates

All genomic data and coordinates are based on the December 2013 version of the genome: hg38, GRCh38.

## 3. RESULTS

### 3.1 Annotation of cell types and subtypes in the human temporal cortex (TC) of individuals with AD, DLB, PD, and neurologically normal controls

Nuclei were isolated from frozen post-mortem human TC tissues of 12 NC donor individuals with no NDD diagnosis or pathological signs, and 12 donors each with diagnoses and corresponding postmortem pathology of AD, DLB, and PD. Each diagnosis group comprised 6 females and 6 males (Table S1 summarizes the demographic and neuropathological phenotypes). snRNA-seq was carried out on prepared gene expression libraries. After quality control (QC) filtering, expression data for nuclei from all four diagnosis groups were integrated, and data from 396,867 nuclei were retained across all four groups (Table S2). Nuclei were then annotated according to major brain cell types by label transfer^34^ from a pre-annotated reference snRNA-seq dataset^35^. These included 19,962 astrocytes, 92,322 excitatory neurons, 44,807 inhibitory neurons, 25,926 microglia, 196,448 oligodendrocytes, and 17,402 oligodendrocyte precursor cells (OPCs). Other cell types including endothelial cells and vascular and leptomeningeal cells made up less than 1% of the total cell population and were therefore excluded from the dataset in downstream analyses.

### 3.2 Vulnerable neuronal types depleted in NDDs compared to neurologically normal controls

AD, DLB, and PD are characterized by the progressive loss of neurons in the brain. To characterize the specific neuronal types that are vulnerable in the temporal cortex of each pathology we performed a comparison analysis of cell-type proportions for each NDD vs NC, restricted to nuclei annotated as excitatory or inhibitory neuronal cells. Expression data for neuronal NC cells were separately integrated with neurons of each NDD. Integrated neuronal cells were then divided into numbered cell subtype clusters, with 30 neuronal subtype clusters for AD, 29 clusters for DLB, and 26 clusters for PD (Fig. 2A). Examination of expression of markers for specific neurotransmitter types among neuronal cell types of each NDD showed the presence of only glutamatergic cell types among excitatory neuron clusters, and GABAergic cell types among inhibitory neuron clusters (Fig. S1A). We then performed a depletion analysis using a beta regression model and calculated the proportion of nuclei from a particular donor sample within each neuronal subtype cluster compared to the total neuronal nuclei for the same sample, and compared the proportions between NDD and NC donors. The results identified four vulnerable neuronal subtypes significantly depleted across the three NDDs (Fig. 2A). Two of these were identified in AD, including one excitatory neuron subtype, AD-Exc7 (*p_adj_*=6.46e-5), and one inhibitory neuron subtype, AD-Inh10 (*p_adj_*=1.90e-5). In DLB, we identified one depleted inhibitory neuron subtype, DLB-Inh10 (*p_adj_*=1.65e-15), and in PD one depleted inhibitory neuron subtype, PD-Inh6 (*p_adj_*=8.53e-7). Of note, the analysis demonstrated that the same inhibitory neuron subtype is depleted in both AD and DLB.

**Figure 2.**
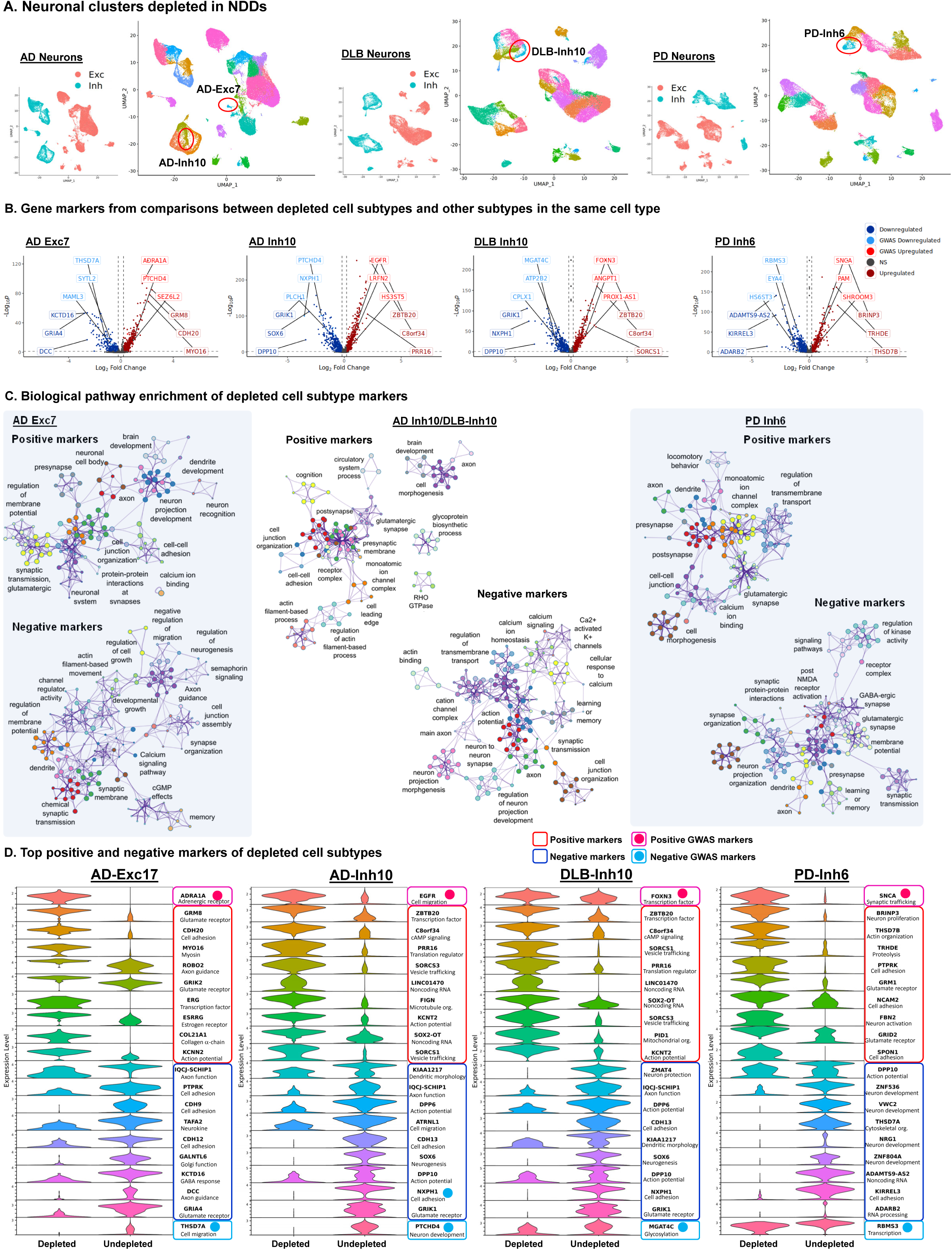
Characterization of vulnerable depleted cell subtypes in each NDD. A. UMAP dimensional reduction plots of neuronal nuclei of each NDD integrated with NC nuclei. Smaller plots are color coded to indicate excitatory neurons (Exc) and inhibitory neurons (Inh). Larger plots are color coded to indicate cell subtype clusters. Depleted clusters are circled in red and labeled. B. Unbiased volcano plots for depleted cell subtype clusters. Log2 fold change (FC) between depleted cluster nuclei and other nuclei of the same major cell type is plotted against – log10 p-value (FDR). Points representing DEGs with statistically significant (FDR < 0.05) upregulation in NDD are shown in dark red while DEGs with significant downregulation are shown in dark blue. Genes without significantly differential expression are shown as gray points. The three DEGs with the highest absolute fold change (log2FC > 0.2) in the up- and downregulated categories are labeled in dark red and dark blue, respectively. The three DEGs within 500kb of NDD-associated SNPs previously identified in GWAS (GWAS-DEG) with the highest absolute log2FC in the up- and downregulated categories are labeled in bright red and bright blue, respectively. C. Metascape network plots of biological pathways enriched among genes upregulated (positive markers) and downregulated (negative markers) within depleted cell subtypes compared to cell subtypes of the same major cell type that were not depleted. Nodes represent specific biological pathways clustered by shared gene membership. Clusters with similar biological function are color coded and labeled according to general function. Node sizes are proportional to the number of differential-interacting genes in the pathway, and line width connecting nodes is proportional to shared gene membership in linked pathways. D. Violin plots of log-normalized count data showing expression of the GWAS-DEGs (bordered in pink and light blue) and 9 overall DEGs (bordered in red and dark blue) with the with the highest absolute fold change in depleted clusters compared to clusters of the same major cell type that were not depleted. Basic functional category information is indicated for each gene.

To characterize the unique transcriptional patterns in the context of disease of each of these depleted subtypes compared to subtypes that were not depleted, we used a likelihood-ratio test to identify differentially expressed genes (DEGs) between each depleted cluster and the other clusters of the same annotated cell type (i.e. excitatory or inhibitory neurons), adjusting for the latent variables age, sex, postmortem interval (PMI), and the number of nuclei after filtering. The comparison was made between the depleted neuronal subtypes and non-depleted subtypes in the disease samples only. DEGs (false discovery rate (FDR) adjusted *p*-value < 0.05) were further categorized into positive (upregulated) and negative (downregulated) genes based on their average log_2_ fold change (Fig. 2B, Tables S3-S6). Strikingly, comparison of positive and negative marker genes across all three depleted inhibitory neuron clusters revealed more than 97% marker gene identity between clusters AD-Inh10 and DLB-Inh10. Furthermore, cell barcode comparison revealed that over 99% of the same NC neuronal cells were present in both clusters, strongly indicating that the two clusters represent the same neuronal subtype, depleted in both AD and DLB. Examination of expression of canonical inhibitory neuron markers used in previous studies^36–40^ among inhibitory subtypes of all NDDs showed the depleted Inh clusters of AD and DLB to be distinguished from other subtypes by strong co-expression of *VIP*, *TAC3*, *PROX1*, *CNR1*, and *TSHZ2*, as well as low expression of *STXBP6*, *LHX6*, *CUX2*, and *PHACTR2*, among other marker genes (Fig. S1B). In contrast, no cell type with a comparable canonical marker expression signature was identified among PD inhibitory neuron clusters.

In order to better understand the biological significance of differential gene expression in the vulnerable neuronal clusters, we examined enrichment of particular biological pathways among positive and negative markers of each depleted subtype^41^, and generated networks of enriched pathways grouped by shared gene membership (Fig. 2C). For all depleted clusters, we primarily found common DEGs associated with functional categories relating to neuronal development and organization (e.g. neuron projection development, axon guidance), synaptic structure (e.g. presynapse, postsynapse, cell-cell adhesion) and synaptic transmission (e.g. regulation of membrane potential, monoatomic ion channel complex, synaptic protein-protein interactions), suggesting that nuances of neuron organization and synaptic function play an important role in determining susceptibility to neurodegeneration.

Examining specific positive and negative marker genes with the most strongly altered (largest fold-change) gene expression in vulnerable neuronal subtypes (Fig. 2D), we found that in AD-Exc7, glutamate receptor-encoding genes *GRM8* and *GRIK2* were among the most strongly upregulated, while the glutamate receptor gene *GRIA4* was among the most strongly downregulated. The cadherin-encoding gene *CDH20*, regulating cell-cell adhesion, was also strongly upregulated, while the cadherin genes *CDH9* and *CDH12* were downregulated, as was *PTPRK*, also involved in cell adhesion. In order to identify marker genes more likely to be involved in driving NDD pathology, we defined genes proximal (within 500Kb) to GWAS-identified risk loci for a particular NDD as “GWAS genes”. Based on GWAS-identified risk loci for AD^10,11^, the adrenergic receptor gene *ADR1A* was the most strongly upregulated AD-GWAS gene marker for AD-Exc7, while the cell migration regulatory gene *THSD7A* was the most strongly downregulated AD-GWAS gene marker.

As noted, depleted subtypes AD-Inh10 and DLB-Inh10 largely shared the same marker genes. The strongest positive markers for both these types included the transcription factor (TF) gene *ZBTB20*, translational regulator *PRR16*, and *SORCS1* and *SORCS3*, both involved in vesicle trafficking and likely playing a role in synaptic transmission. The most strongly upregulated AD-GWAS gene marker was *EGFR*, involved in cell migration, while the most strongly downregulated AD-GWAS gene marker was *PTCHD4*, involved in neuronal development. Based on GWAS-identified risk loci for DLB^15,16^, the most strongly upregulated DLB-GWAS gene marker was the TF-encoding *FOXN3*, while the most strongly downregulated DLB-GWAS gene was *MGAT4C*, involved in protein glycosylation.

The subtype depleted in PD, PD-Inh6, showed marked upregulation of glutamate receptor genes *GRM1* and *GRID2*, as well as cell adhesion-regulating genes *NCAM2* and *SPON1*, while downregulation of several developmental genes was observed, including *ZNF536*, *VWC2*, *NRG1*, and *ZNF804A*. Notably, the most strongly upregulated PD-GWAS gene marker (based on GWAS-identified risk loci for PD^13^) for this cluster was *SNCA*, suggesting that overexpression of the *SNCA* gene correlates with vulnerability to neurodegeneration in PD. The most strongly downregulated PD-GWAS gene marker was the transcriptional regulatory gene *RBMS3*.

### 3.3 Characterization of disease-driver cell subtypes with enriched expression of GWAS-identified risk genes

We sought to identify cell subtypes that were potentially important for conferring risk of each NDD, hereafter disease-driver cell types, based on increased expression of GWAS genes. First, we integrated, annotated, and clustered nuclei of each NDD with NC nuclei as described above, except that in this case nuclei of all cell types, including astrocytes (Astro), excitatory neurons (Exc), inhibitory neurons (Inh), microglia (Micro), oligodendrocytes (Oligo), and oligodendrocyte precursor cells (OPC) were included rather than neuronal nuclei alone. This resulted in delineation of 32 cell subtype clusters in AD, 32 clusters in DLB, and 35 clusters in PD (Fig. 3A). We next examined each subtype for enriched expression of GWAS genes using AUCell^42^. This program compares expression of a defined gene set (i.e. GWAS proximate genes) to total genes expressed in each nucleus, and determines whether the gene set is expressed in a significantly higher proportion than would be expected by chance. We defined a cluster as a disease-driver if over 99% of nuclei showed significant enrichment for GWAS gene set expression. In this way we identified one disease-driver oligodendrocyte cluster in AD (AD-Oligo3), four disease-driver excitatory neuron clusters (DLB-Exc1, 5, 8, 10) and two inhibitory neuron clusters (DLB-Inh1, 2) in DLB, and four disease-driver excitatory neuron clusters in PD (PD-Exc4, 5, 6, 7) (Fig. 3A, B). Thus, both DLB and PD produced multiple neuronal cell types that were implicated as disease drivers, while in AD only a single oligodendrocyte disease-driver cell subtype was identified.

**Figure 3.**
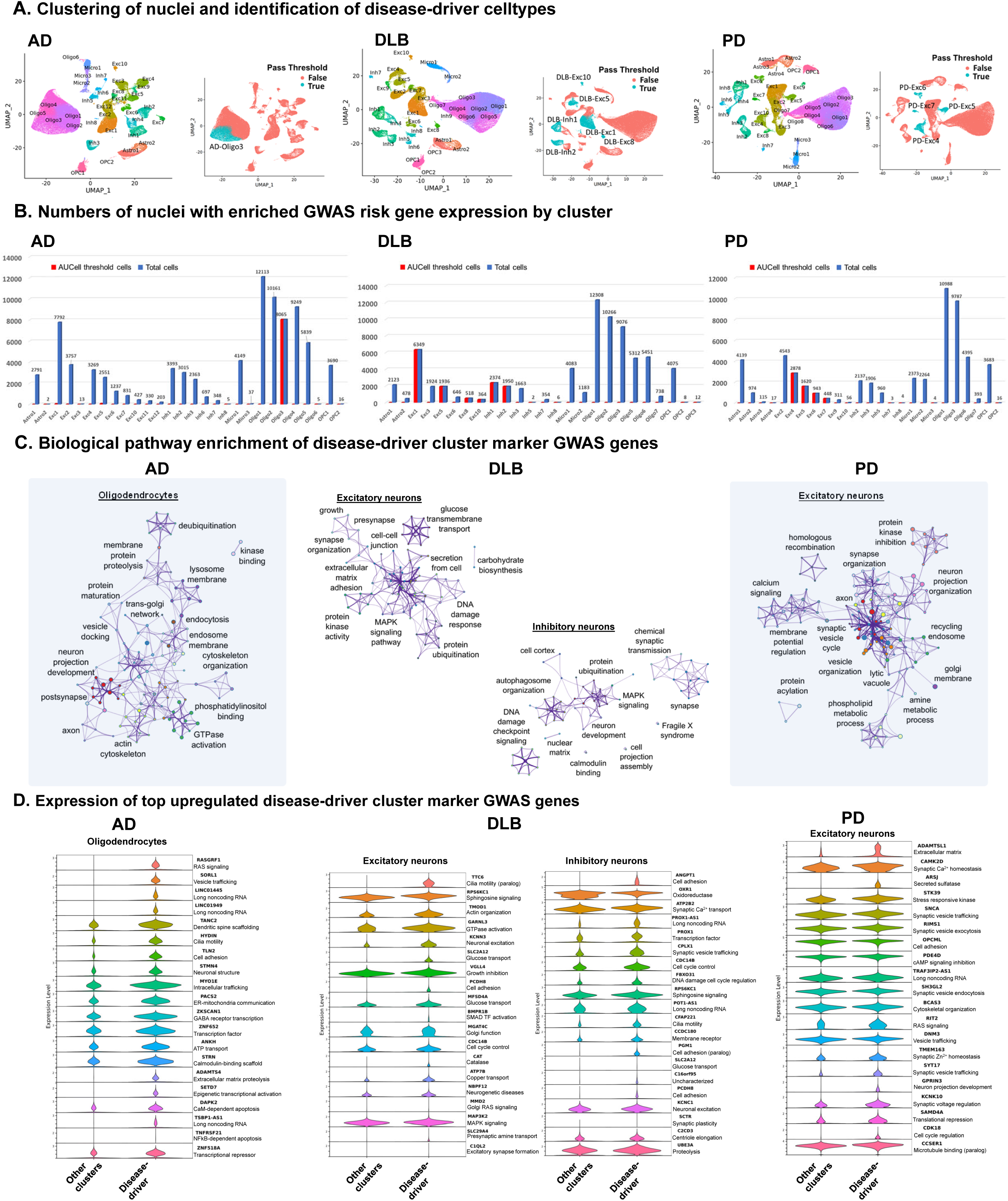
Identification of disease-driver cell subtypes with enriched GWAS risk gene expression. A. UMAP dimensional reduction plots of neuronal nuclei of each NDD integrated with NC nuclei. Smaller plots are color coded to indicate subtypes below (False) and above (True) the AUCell pass threshold for enriched expression of genes within 500kb of NDD-associated SNPs previously identified in GWAS (GWAS genes). B. Bar charts showing total numbers of cells in each subtype cluster (blue) and numbers of cells above the AUCell pass threshold for enriched GWAS gene expression (red). C. Metascape network plots of biological pathways enriched among GWAS genes upregulated within disease-driver cell subtypes compared to cell subtypes of the same major cell type that were enriched for GWAS gene expression. Nodes represent specific biological pathways clustered by shared gene membership. Clusters with similar biological function are color coded and labeled according to general function. Node sizes are proportional to the number of differential-interacting genes in the pathway, and line width connecting nodes is proportional to shared gene membership in linked pathways. D. Violin plots of log-normalized count data showing expression of the GWAS-DEGs with the highest positive fold change in disease-driver clusters compared to clusters of the same major cell type that were not disease-driving. Basic functional category information is indicated for each gene.

In order to understand the potential functional significance of risk genes expressed in these disease-driver clusters, we performed marker gene analysis as above, comparing gene expression in disease-driver clusters of a particular cell type to all of the other clusters of that same cell type in NDD nuclei (Tables S7-S10). We then examined biological pathway enrichment among GWAS genes upregulated in each set of disease-driver cell types. Finally, we clustered enriched pathways based on common gene membership (Fig. 3C). In the disease-driver oligodendrocyte cluster of AD, AD-Oligo3, we found enrichment of numerous pathways relating to endosomal vesicle trafficking (specific strongly upregulated genes relating to this pathway including *SORL1*, *MYO1E*, and *PACS2* (Fig. 3D)), cytoskeletal organization (e.g. *HYDIN*, *TANC2*, *STRN*), and regulation of proteolysis (e.g. *ADAMTS4*) and apoptosis (e.g. *DAPK2*, *TNFRSF21*). Notably, we also observed strongly inhibited expression of the major AD risk factor gene *BIN1* in this cell type (Table S7). In disease-driver excitatory neuron clusters of DLB, we identified enrichment of pathways relating to synaptic organization and transmission (e.g. *KCNN3*, *SLC29A4*, *C1QL2*), cell adhesion (e.g. *PCDH8*), transmembrane transport (e.g. *SLC2A12*, *MSFD4A*, *ATP7B*), DNA damage response (e.g. *CDC14B*), and proteolysis. Among disease-driver inhibitory neurons in DLB, we found enrichment of pathways relating to synaptic transmission (e.g. *ATP2B2*, *CPLX1*, *KCNC1*, *SCTR*), autophagy, proteolysis (e.g. *UBE3A*), and DNA damage response (e.g. *CDC148*, *FBXO31*). In disease-driver excitatory neurons of PD, we found enrichment of risk genes involved in synaptic organization and transmission (e.g. *SNCA*, *CAMK2D*, *RIMS1*, *SH3GL2*, *TMEM163*, *SYT17*, *KCNK10*), autophagy, phospholipid metabolism, and homologous recombination. It is notable that as for the PD-depleted neuron cluster above, *SNCA* was also among the top upregulated GWAS genes within PD-disease driver neuron clusters.

### 3.4 Altered cell to cell communication pathways in NDDs

Next, we aimed to investigate changes in interactions between different cellular subtypes associated with each of the three NDDs. To accomplish this, we used the same integrated datasets of NC nuclei and nuclei of each NDD used above for analysis of disease-driver subtypes. We analyzed expression of known interacting ligands and receptors in each of the subtype clusters to identify pairs of subtypes with likely communication using CellChat^43^. Predicted interactions were then compared between NC and NDD nuclei to identify disease-associated changes in cell-cell communication. Comparisons were made with regard to relative strength of interactions between cell subtypes based on changes in gene expression levels between NC and NDD nuclei of the same subtype.

Changes in interaction strength were varied across the three NDDs (Fig. 4A). In AD, such changes were overall split between increased and decreased communication among different cell types, with both large increases and decreases observed among the top 10% of altered cell type interactions. The cell types with the largest increases in interaction strength included several excitatory neuron subtypes, AD-Exc1, 3, and 4, and inhibitory neuron subtype AD-Inh1, as well as oligodendrocyte subtypes AD-Oligo1 and 4. All of these cell types showed primarily increased communication with neuronal subtypes. In contrast, decreased interaction strength was observed in astrocyte cluster AD-Astro1, excitatory neuron cluster AD-Exc2, and oligodendrocyte precursor cell cluster AD-OPC1, all of which showed reduced communication with one another as well as with several neuronal and oligodendrocyte subtypes. In DLB, by contrast, overall changes primarily showed decreases in interaction strength. Among the strongest effects, subtypes DLB-Astro1, DLB-Exc1, 3, 5, and 6, DLB-Inh1, 2, 3, and 4, DLB-Oligo1 and 5, and DLB-OPC1 showed reduced communication strength mainly with one another. However, subtypes DLB-Oligo1, 2, 3, 4, and 6 showed increased communication with one another as well. In PD, overall decreased interaction strength was also observed, with the strongest decreases found between the cell types PD-Astro1 and 2, PD-Exc1, 2, 3, 5, and 6, PD-Inh2, and 4, PD-Oligo1, and PD-OPC1. Increased interaction strength in PD was observed for clusters PD-Oligo2, and 4, primarily with regard to other oligodendrocyte clusters. Overall the results demonstrated increased interaction strength in AD driven primarily by excitatory neurons and oligodendrocytes, but decreased interaction strength in DLB and PD, driven primarily by both inhibitory and excitatory neurons, as well as oligodendrocytes. Thus, changes in cell-cell communication strength in DLB and PD closely resembled one another, while patterns in AD were more distinct.

**Figure 4.**
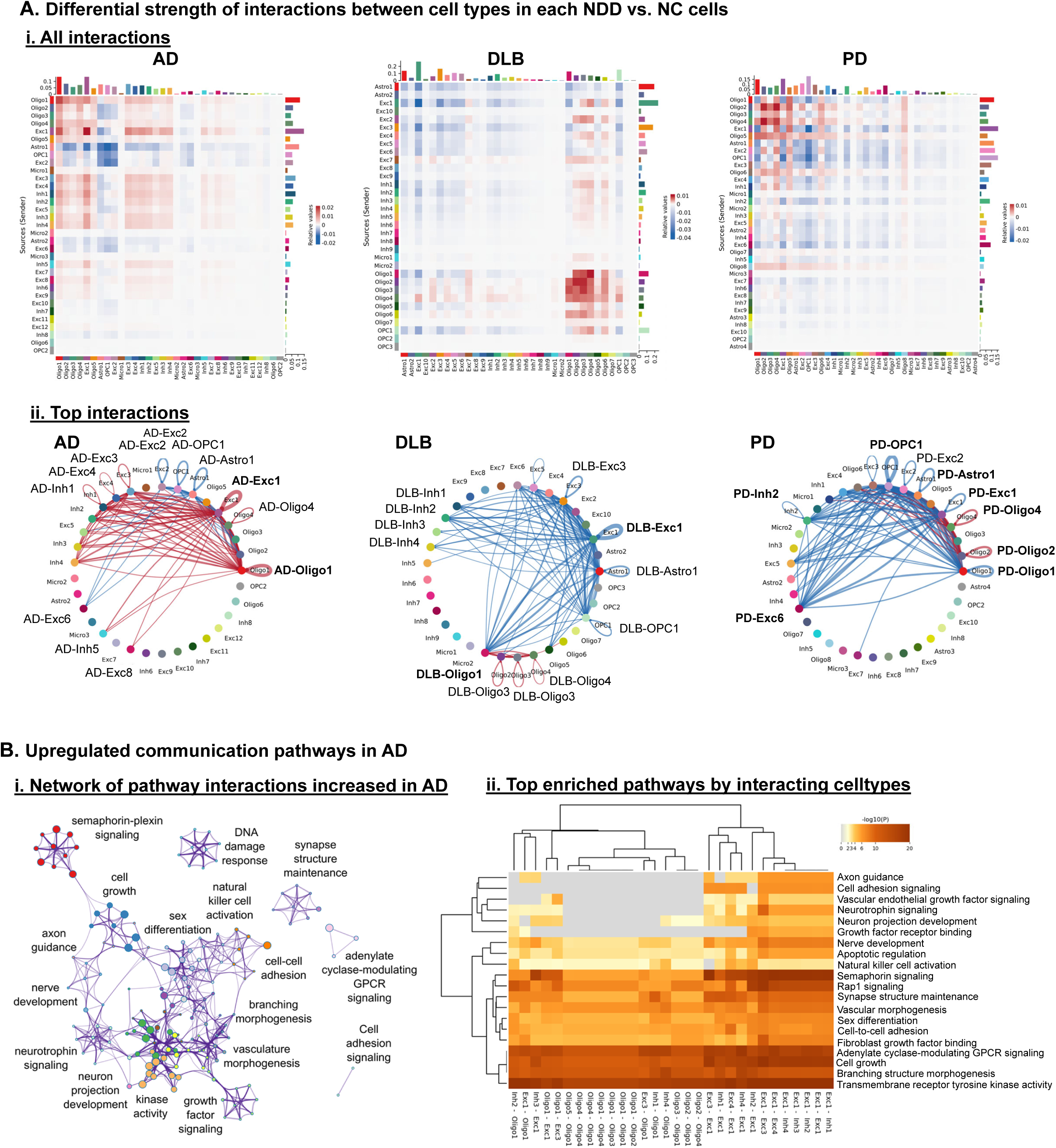
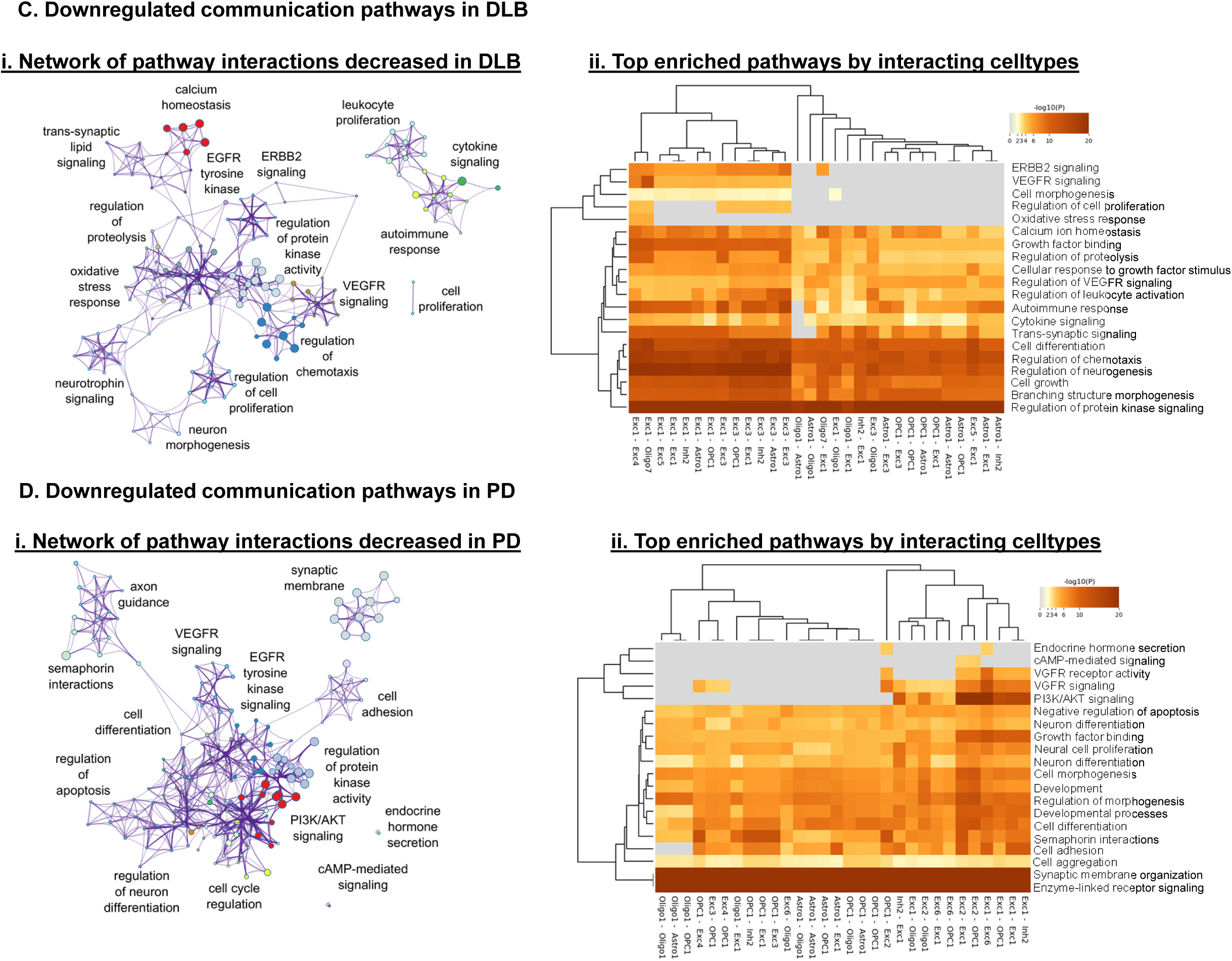
Differential interaction strength between cell subtypes in NDDs vs. Normal nuclei. A. *i.* CellChat heatmaps showing degree of overall change in interaction strength between all pairs of cell subtypes for each NDD. Red indicates increased interaction in NDD, blue indicates decreased interaction. *ii.* CellChat network diagram showing celltypes with the highest differential interaction strength based on fold change in receptor-ligand expression in NDD nuclei compared to NC. Lines between celltypes indicate significantly altered interaction, with red lines indicating increased interaction strength in NDD and blue lines representing decreased interaction strength. Line width is proportional to statistical significance of change in interaction strength. Larger and bold labels indicate celltypes with more prominently altered interactions. B. *i.* Metascape network plot of biological pathways enriched among genes associated with increased interaction strength in AD across all celltypes. Nodes represent specific biological pathways clustered by shared gene membership. Clusters with similar biological function are color coded and labeled according to general function. Node sizes are proportional to the number of differential-interacting genes in the pathway, and line width connecting nodes is proportional to shared gene membership in linked pathways. *ii.* Heatmap of top 20 enriched pathways among interactions increased in AD across all celltypes. Interacting celltypes are indicated, with sending type listed first and receiving type indicated second. Color saturation is proportional to strength of enrichment. C. *i.* Metascape network plot of biological pathways enriched among genes associated with increased interaction strength in DLB across all celltypes. *ii.* Heatmap of top 20 enriched pathways among interactions increased in DLB across all celltypes. D. *i.* Metascape network plot of biological pathways enriched among genes associated with increased interaction strength in PD across all celltypes. *ii.* Heatmap of top 20 enriched pathways among interactions increased in PD across all celltypes.

To get new insights into the biological significance of cell-cell communication in the three NDDs, we examined the biological pathway associations of the genes involved in altered communication between each pair of cell subtypes using Metascape. Pathways enriched among genes associated with the top AD-increased interactions related primarily to cell growth, development, and morphology, as well as DNA damage response, stress response, and GPCR and kinase signaling (Fig. 4Bi). The pathways enriched among AD-increased interactions across all cell types notably differed between neuron-to-neuron interactions and oligodendrocyte-to-neuron interactions (Fig. 4Bii). Pathways strongly enriched among all interaction types were associated with cell growth and morphogenesis, and GPCR and tyrosine kinase receptor signaling, while interactions more strongly enriched in neuron-to-neuron interactions related specifically to nerve morphogenesis and organization, including axon guidance, nerve development, semaphorin signaling, and neurotrophin signaling.

In DLB, interaction strength was overall reduced compared to NC nuclei, and pathways enriched among genes associated with the top DLB-decreased interactions related primarily to cell growth and development, immune response signaling, and calcium homeostasis (Fig. 4Ci). Pathway enrichment was strongest in DLB-decreased communications involving the Exc1 and Exc3 excitatory neuron subtypes as the transmitting cell type, with a wide variety of receiving cell types (Fig. 4Cii). Pathways enriched specifically in these types of interactions related to cell growth and proliferation, cell morphogenesis, and the oxidative stress response. Pathways enriched among all interacting cell types additionally included calcium ion homeostasis, immune response signaling, chemotaxis, proteolysis, and general kinase signaling.

In PD, interaction strength was also reduced overall. Pathways enriched among genes associated with the top PD-decreased interactions again related to cell growth and development, and also to axon guidance and neuronal organization, synaptic membrane structure, and regulation of apoptosis (Fig. 4Di). Some specific pathways were most often enriched in PD-decreased communications in which neuronal subtypes were the transmitting cell type, including PI3K/AKT growth signaling, cAMP signaling, and endocrine hormone signaling (Fig. 4Dii). Many pathways involved in growth and development were enriched across all interaction types, as were pathways associated with regulation of apoptosis, cell adhesion, synaptic membrane organization, and enzyme-linked receptor signaling.

Next, to organize altered cell-to-cell communication networks with regard to the specific cell types involved, individual pairs of interacting proteins in NDD and NC nuclei were grouped by association with particular biological pathways, and each of these pathway groups were further clustered based on the particular cell subtypes in communication, following principal component analysis (PCA) (Fig. S2A). This led to the identification of four communication clusters each in AD and DLB, and five clusters in PD. In AD and PD, each cluster contained a qualitatively even distribution of pathways from both NC and NDD nuclei. However, in DLB, cluster 1 was entirely composed of communication pathways identified in NC nuclei, while cluster 3 was heavily dominated by pathways identified in DLB nuclei, suggesting the development of distinct cell-to-cell communication networks in the context of DLB (Fig. S2B).

### 3.5 Shared patterns of differential gene expression among NDDs

In order to identify commonalities in gene dysregulation among NDDs, we integrated snRNA-seq data from nuclei of all three NDDs and NC nuclei for each of the six major cell types and grouped these into cell subtype clusters as described above. Next we further annotated these clusters as more specific predicted cell types using the scMayoMap^44^ software package (Fig. 5A), and employed the NEBULA^45^ software package to perform differential gene expression analysis between NC nuclei and those of each NDD at the cell subtype level. Across all three NDDs, the highest numbers of DEGs were identified in inhibitory neuron subtypes, and the majority were downregulated (Fig. 5B). Most excitatory neurons and astrocytes clusters in AD exhibited primary gene downregulation, while, in DLB and PD both upregulated and downregulated DEGs were detected in those clusters. On the other side, microglia showed mixed up- and downregulation in AD, but predominantly upregulation in DLB and PD in most subtypes. OPC subtypes showed both up- and downregulation DEGs within each NDD. Oligodendrocytes were also varied, with mixed distribution of up- and downregulation in AD, predominant upregulation in DLB, and predominant downregulation in PD. Notably, *SNCA* was upregulated in DLB in four separate oligodendrocyte clusters (Oligodendrocyte 1, 3, 5, and 10), but not in oligodendrocyte clusters of PD, suggesting a potentially important function in oligodendrocytes for this key synucleopathy gene specifically in the context of DLB.

**Figure 5.**
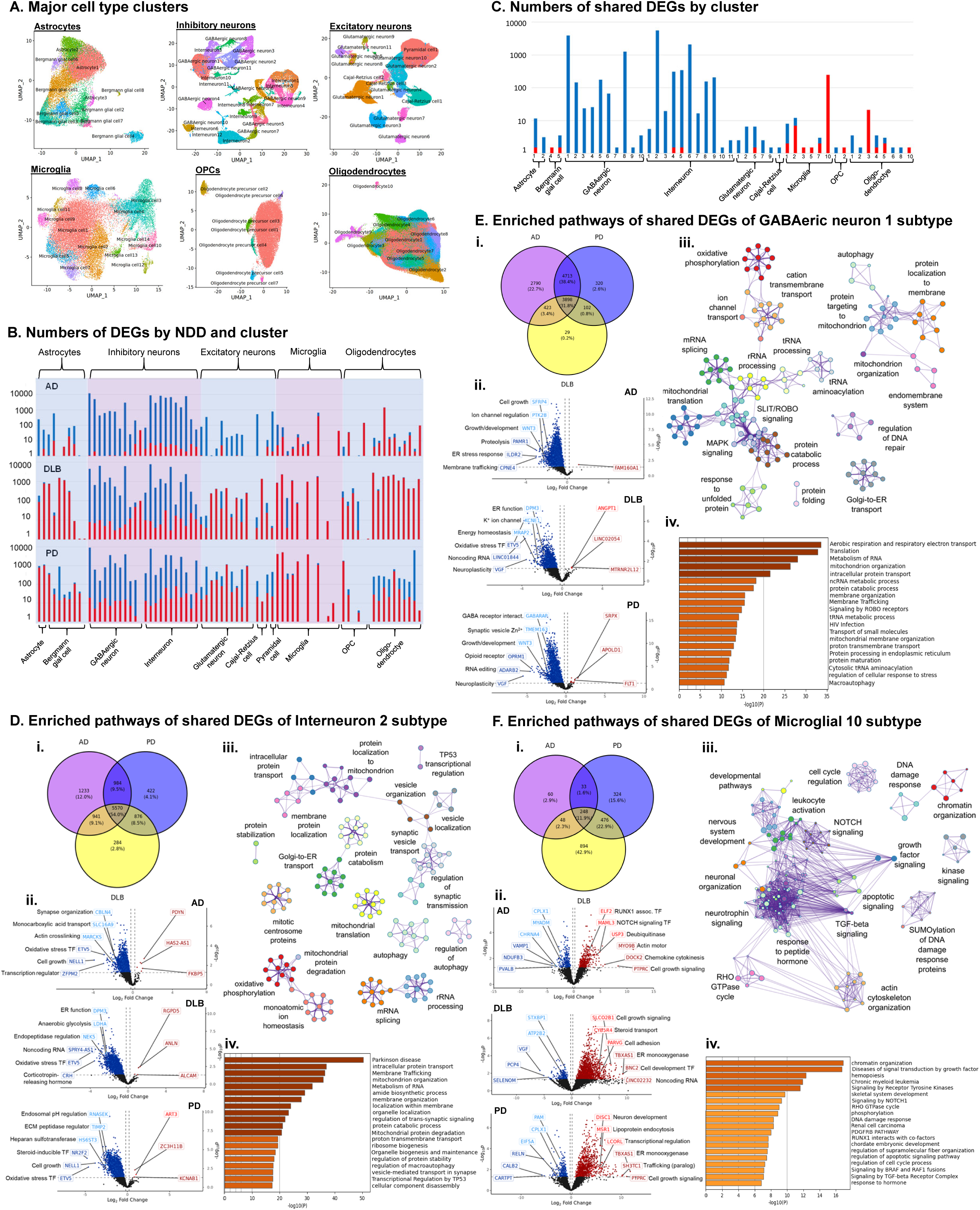
Differential gene expression shared by three pathologies on cell subtype level. A. UMAP dimensional reduction plots of integrated NDD and NC nuclei of each major cell type, color coded to indicate cell subtype clusters. B. Bar charts representing numbers of DEGs identified in each cell subtype within each NDD compared to NC nuclei of the same subtype. Red indicates DEGs upregulated in NDDs and blue indicates DEGs downregulated in NDDs. C. Bar chart representing numbers of DEGs shared between all 3 NDDs compared to NC nuclei for each cell subtype. Red indicates DEGs upregulated in NDDs and blue indicates DEGs downregulated in NDDs. D. *i.* Venn diagram showing overlap between DEGs downregulated in each NDD within the Interneuron 2 subtype. *ii.* Unbiased volcano plots for GABAergic neuron 1 subtype gene expression in each NDD. Log2 fold change (FC) between NDD nuclei and NC nuclei of the same subtype is plotted against –log10 p-value (FDR). Points representing DEGs with statistically significant (FDR < 0.05) upregulation in NDD are shown in dark red while DEGs with significant downregulation are shown in dark blue. Genes without significantly differential expression are shown as gray points. The three DEGs with the highest absolute fold change (log2FC > 0.2) in the up- and downregulated categories are labeled in dark red and dark blue, respectively. The three DEGs within 500kb of NDD-associated SNPs previously identified in GWAS (GWAS-DEG) with the highest absolute log2FC in the up- and downregulated categories are labeled in bright red and bright blue, respectively. Basic functional category information is indicated for each labeled GWAS-DEG. *iii.* Metascape network plots of biological pathways enriched among DEGs downregulated in all NDDs within the GABAergic neuron 1 subtype. Nodes represent specific biological pathways clustered by shared gene membership. Clusters with similar biological function are color coded and labeled according to general function. Node sizes are proportional to the number of differential-interacting genes in the pathway, and line width connecting nodes is proportional to shared gene membership in linked pathways. *iv.* Metascape bar chart showing the top 20 most highly enriched biological pathway terms among DEGs downregulated across all NDDs within the GABAergic neuron 1 subtype. Statistical significance (Log10 p-value) is plotted on horizontal axes. Darker-colored bars indicated greater significance. E. *i.* Venn diagram showing overlap between DEGs downregulated in each NDD within the GABAergic neuron 1 subtype. *ii.* Unbiased volcano plots for Interneuron 2 subtype gene expression in each NDD. *iii.* Metascape network plots of biological pathways enriched among DEGs upregulated in all NDDs within the Interneuron 2 subtype. *iv.* Metascape bar chart showing the top 20 most highly enriched biological pathway terms among DEGs downregulated across all NDDs within the Interneuron 2 subtype. F. *i.* Venn diagram showing overlap between DEGs upregulated in each NDD within the Microglia 10 subtype. *ii.* Unbiased volcano plots for Microglia 10 subtype gene expression in each NDD. *iii.* Metascape network plots of biological pathways enriched among DEGs upregulated in all NDDs within the Microglia 10 subtype. *iv.* Metascape bar chart showing the top 20 most highly enriched biological pathway terms among DEGs upregulated across all NDDs within the Microglia 10 subtype.

Next, for each cell subtype we catalogued the shared up- and downregulated DEGs across all three NDDs (Fig. 5C). As expected, inhibitory neuron subtypes exhibited the highest number of DEGs and almost all were downregulated. The Interneuron 2 inhibitory neuron subtype exhibited the highest number of shared downregulated DEGs (5,570; Fig. 5D, Table S10). followed by the GABAergic neuron 1 subtype (3,898; Fig. 5E, Table S11). Additionally, about 900 downregulated DEGs were shared between each pair of pathologies in Interneuron 2 (984 for AD and PD, 941 for AD and DLB, 876 for DLB and PD; Fig. 5Di). Similarly, GABAergic neuron 1 also exhibited additional shared DEGs between each pair of NDDs (4,713 for AD and PD, 423 for AD and DLB, 102 for DLB and PD; Fig. 5Ei). Microglia 10 had the highest number of shared upregulated DEGs (248; Fig. 5F, Table S12). Examination of overlap between each pair of pathologies in Microglia 10 identified the largest number of shared upregulated DEGs (476) between DLB and PD, and fewer shared DEGs between the other pairs (48 for AD and DLB, 33 for AD and PD; Fig. 5Fi). In contrast, other major cell types shared only a relatively small number of DEGs. Overall, these results suggested that the common dysregulated pathways across NDDs are mainly found in inhibitory neurons.

Thus, we next analyzed the enrichment of biological pathways among shared downregulated DEGs in the Interneuron 2 and GABAergic neuron 1 subtypes. As these are pathways enriched among downregulated DEGs they may reflect impaired biological pathways. In the Interneuron 2 subtype, we identified enrichment of pathways related to synaptic vesicle transport, mitochondrial function, oxidative phosphorylation, autophagy, proteolysis, and RNA processing (Fig. 5Dii). These functional categories were also identified in the analysis of the top enriched individual pathways (Fig. 5Diii). Specific genes that were strongly downregulated in all three NDDs included the transcription factor (TF) gene *ETV5*, associated with the response to oxidative stress, and the cell growth regulator gene *NELL1*, as well as the AD-GWAS gene *CBLN4*, involved in synapse organization, the DLB- and PD-GWAS gene *DPM3*, involved in endoplasmic reticulum (ER) function, and the autophagy-associated PD-GWAS gene *RNASEK* (Fig. 5Div). The respective DLB- and PD-GWAS genes *NEK5* and *TIMP2*, both involved in regulation of proteolysis, were strongly downregulated in both DLB and PD.

In the GABAergic neuron 1 subtype, the identified enriched pathways based on shared downregulated DEGs were overall similar to those of Interneuron 2 (Fig. 5Eii), including aerobic respiration and respiratory electron transport, translation, metabolism of RNA, and mitochondrion organization (Fig. 5Eiii). *ETV5* and *DPM3* were again among the most highly downregulated genes in all three NDDs, as was the AD-GWAS gene *VGF*, involved in regulation of neuroplasticity, and the AD- and PD-GWAS GABA-receptor interacting gene *GABARAP* (Fig. 5Eiv). Developmental regulator *WNT3*, a GWAS gene for both AD and PD, was also highly downregulated in those two NDDs.

Similarly, we analyzed pathway enrichment in upregulated DEGs of the Microglia 10 subtype, plausibly indicating activation of biological pathways. The results demonstrated enrichment for growth and developmental pathways, as well as pathways associated with leukocyte activation, cell cycle regulation, DNA damage response, chromatin organization, and cytoskeletal organization (Fig. 5Fii). The strongest enriched individual pathways included chromatin organization, growth factor signal transduction, receptor tyrosine kinase signaling, and NOTCH1 signaling (Fig. 5Fiii). The TF genes *ELF2* and *MAML3*, and the deubiquitinase gene *USP3*, all AD-GWAS genes, and the transcriptional regulator PD-GWAS gene *LCORL* were among the most strongly overexpressed DEGs across all three NDDs, as were the actin motor gene *MYO9B*, and the cell growth signaling gene *PTPRC* (Fig.4Fiv). The gene *DOCK2*, involved in chemokine-responsive cytokinesis, was strongly upregulated in both AD and DLB, while the DLB-GWAS gene *SLCO2B1*, also involved in cell growth signaling, the steroid transport gene *CYB5R4*, the PD-GWAS gene *DISC1*, regulating neuronal development, and the ER monooxygenase gene *TBXAS1*, were strongly upregulated in both PD and DLB. In summary, we observed high numbers of shared downregulated genes in inhibitory neuron subtypes across all three NDDs, indicating impairment of pathways relating to neuronal development, synaptic function, stress responses, and other categories, but more diverse expression patterns in other types, with fewer shared DEGs.

### 3.6 Differential gene expression between NDDs

To advance the understanding of mechanistic diversity amongst NDDs we next studied the differential transcriptomic landscape between NDDs. To accomplish this, we integrated transcriptomic data for all cell types from each pair of NDDs (i.e., AD and DLB, PD and DLB, and AD and PD) and performed dimensional reduction and clustering of the integrated datasets to identify cell subtypes (Fig. 6A). Differential expression analysis was performed at the cell subtype level for each NDD pairing to identify distinct DEGs between the pathologies. In comparing AD and DLB, we found DEGs that were upregulated in DLB in only four out of the 29 cell subtype clusters, including excitatory neurons (clusters 5 and 9), and oligodendrocytes (clusters 1 and 2), which exhibited about 5,000 DEGs each (5347, 5030, 4630, and 4805, respectively), mainly upregulated in DLB (Fig. 6B, Ci, Tables S13-S16). The only other clusters that exhibited more than 100 DEGs were Exc3 and Oligo6. Biological pathway enrichment analysis of DLB-upregulated DEGs in the excitatory neuron subtypes revealed enrichment of genes involved in cell cycle regulation, synaptic transmission, and stress response. In oligodendrocyte clusters we found enrichment for pathways associated with inclusion body assembly, cellular signaling, and chromatin organization (Fig. 6Cii). In addition, genes involved in DNA damage response, proteolysis, immune response, and transcriptional regulation were enriched in both of these cell types. Accordingly, the strongest DLB-upregulated genes also play roles in these functional categories, including GWAS risk genes for both AD and DLB (Fig. 6Ci). For example, *RTF2,* a DEG in Exc5 and Oligo2, and *FBXO31* in Oligo 2 are involved in DNA damage response, and the DEGs *SUGT1* in Exc5, *CCNE2* in Exc9, and *GAK* in Oligo1 and 2, among others, are involved in cell cycle regulation. The proteolysis associated gene *MAEA* is a GWAS risk gene for both AD and DLB and was among the highest DLB-upregulated DEGs in both Exc9 and Oligo1. The growth factor signaling AD-GWAS gene *PLCG2* was highly DLB- upregulated in all four cell types.

**Figure 6.**
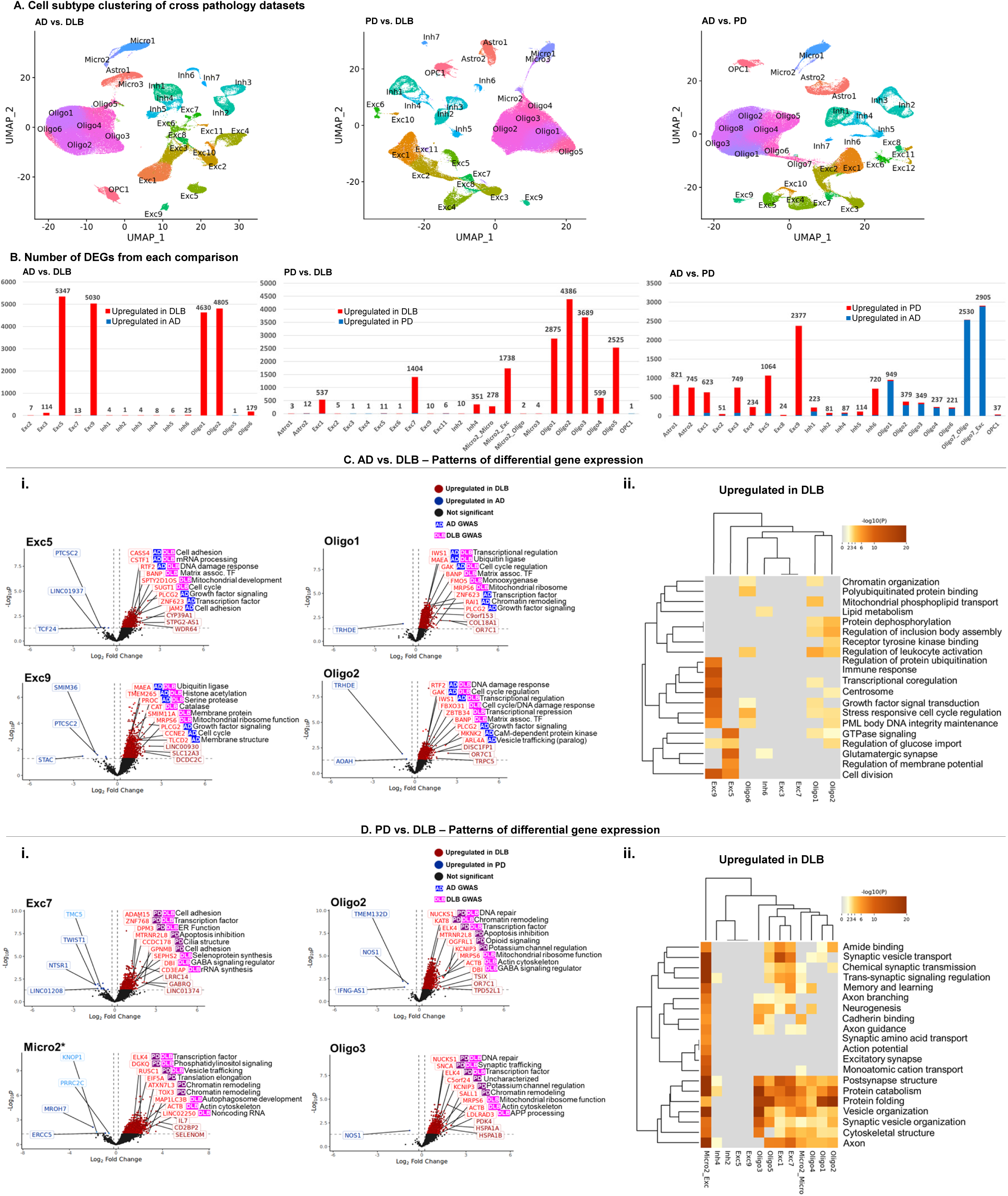
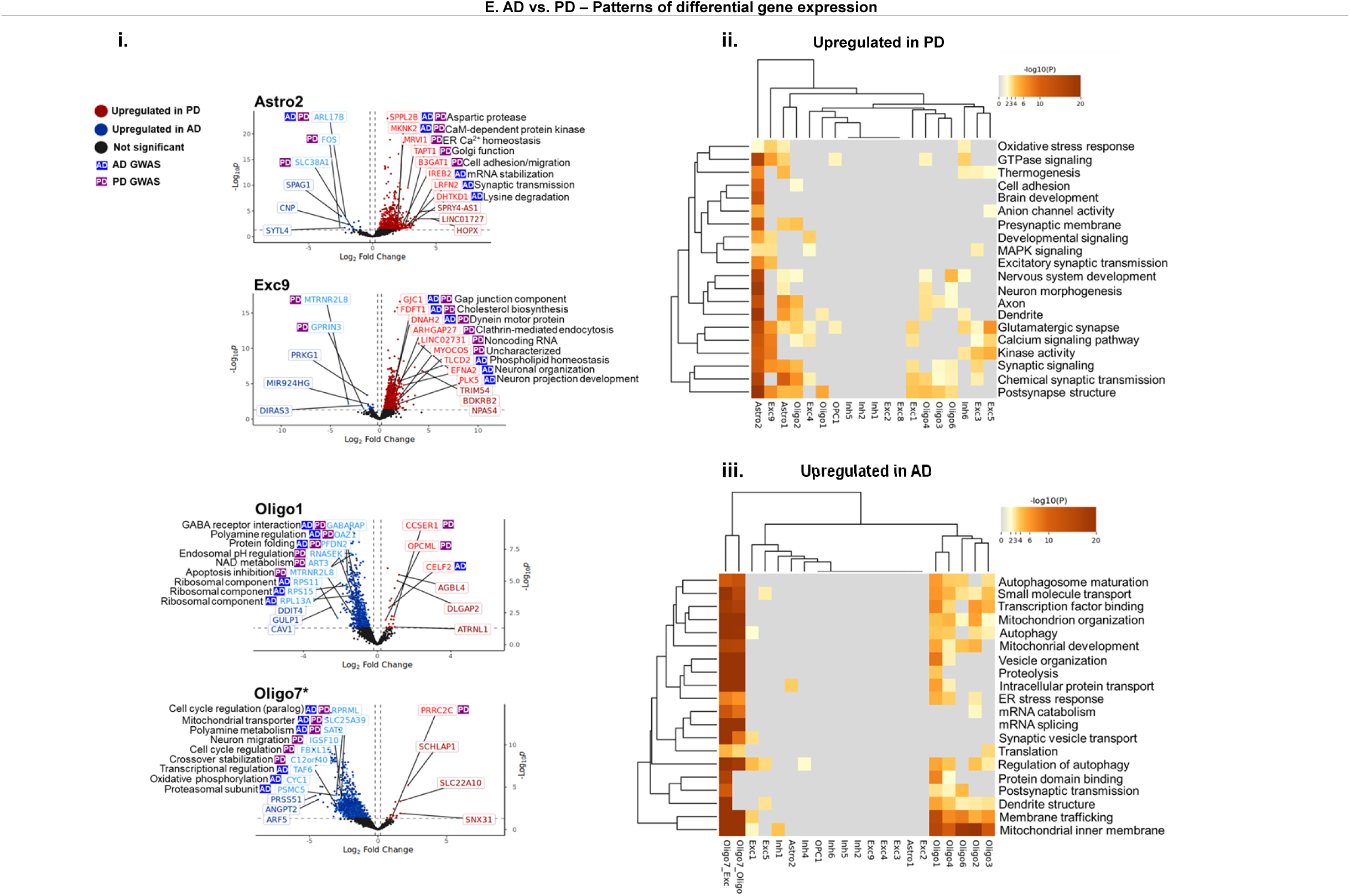
Differential gene expression between NDDs in cell subtypes. A. UMAP dimensional reduction plots of integrated pairs of NDD nuclei of all cell types, color coded to indicate cell subtype clusters. B. Bar charts representing numbers of DEGs identified using NEBULA for each cell subtype between nuclei of the indicated NDD pairs within the same subtype. Red and blue bars represent DEGs upregulated in one or the other NDD, as indicated. C. *i.* Unbiased volcano plots showing gene expression in selected cell subtypes in the AD and DLB comparison. Log2 fold change (FC) between nuclei of the 2 NDDs in the same subtype is plotted against –log10 p-value (FDR). Points representing DEGs with statistically significant (FDR < 0.05) upregulation in AD are shown in dark blue while DEGs with significant upregulation in DLB are shown in dark red. Genes without significantly differential expression are shown as gray points. The three DEGs with the highest absolute fold change (log2FC > 0.2) in the AD and DLB upregulated categories are labeled in dark blue and dark red, respectively. The three DEGs within 500kb of NDD-associated SNPs previously identified in GWAS (GWAS-DEG) exclusive to AD, exclusive to DLB, and common to both NDDs with the highest absolute log2FC in the up- and downregulated categories are labeled in bright red and bright blue, respectively, and the NDDs associated with each GWAS-DEG are indicated. Basic functional category information is indicated for each labeled GWAS-DEG. *ii.* Heatmap of top 20 enriched pathways among interactions increased in DLB compared to AD across all celltypes. Color saturation is proportional to statistical significance of enrichment. D. *i.* Unbiased volcano plots showing gene expression in selected cell subtypes in the PD and DLB comparison. Color coding indicates upregulation in the indicated NDD. The top three GWAS-DEGs exclusive to PD, exclusive to DLB, and common to both NDDs are indicated. *ii.* Heatmap of top 20 enriched pathways among interactions increased in DLB compared to PD across all celltypes. E. *i.* Unbiased volcano plots showing gene expression in selected cell subtypes in the AD and PD comparison. Color coding indicates upregulation in the indicated NDD. The top three GWAS-DEGs exclusive to AD, exclusive to PD, and common to both NDDs are indicated. *ii.* Heatmap of top 20 enriched pathways among interactions increased in PD compared to AD across all celltypes. *iii.* Heatmap of top 20 enriched pathways among interactions increased in AD compared to PD across all celltypes.

Comparison of PD to DLB across all clusters also resulted mainly in DLB-upregulated DEGs (Fig. 6B, Di, Tables S17-S20). Genes were strongly upregulated in DLB in a number of oligodendrocyte clusters (2875, 4386, 3689, and 2525 in Oligo1, 2, 3, and 5, respectively), with fewer DEGs in excitatory neuron clusters (537 and 1404 in Exc1 and 4, respectively). Additionally, while the Micro2 cluster was annotated as a microglial cluster due to this being the most prevalent cell type, excitatory neuron nuclei comprised approximately a third of the cluster and >10% of the cluster was made up of oligodendrocyte cells. For this reason we separately performed differential expression analysis on each of these three cell types within the cluster. We identified 6.25-fold more DEGs for the excitatory neuron subset (Micro2_Exc) compared to the microglial subset (Micro2_Micro), indicating excitatory neurons as the primary source of differential gene expression for this cluster. Biological pathway analysis revealed that the top enriched pathways across cell subtypes included synaptic transmission, neuronal morphology, protein folding and proteolysis (Fig. 6Dii). The strongest enrichment was observed in Micro2 excitatory neurons followed by multiple oligodendrocyte and other excitatory neuron subtypes, as well as Micro2 microglia. Synaptic transmission-associated pathways were most strongly enriched in excitatory neuron subtypes. DLB- and PD-GWAS genes strongly upregulated in DLB were also associated with these functional categories, including synaptic adhesion-related genes *ADAM15* and *GPNMB* in Exc7, and synaptic vesicle-trafficking gene *RUSC1* in Micro2 (Fig. 6Di). Chromatin remodeling GWAS genes were DLB-upregulated across multiple clusters, including *ATXN7L3* and *TOX3* in Micro2, *KAT8* in Oligo2, and *SALL1* in Oligo3, while the TF *ELK4* was DLB-upregulated in all three clusters. The DNA repair-associated gene *NUCKS1* and actin gene *ATCB* were highly DLB-upregulated in both oligodendrocyte clusters Oligo2 and 3. Notably, *SNCA* and the amyloid precursor protein (APP)-processing gene *LDLRAD3* were both among the most highly DLB-upregulated GWAS genes in Oligo3.

Comparing AD to PD yielded the most diverse pattern of transcriptional dysregulation as demonstrated by the variety of cell types with DEGs and the directionality of the differential expression (Fig. 6B, Ei, Tables S21-S25). Upregulation in PD was observed in astrocyte (821 and 745 DEGs in Astro1 and 2, respectively), excitatory (623, 749, 1064, and 2377 in Exc1, 3, 5, and 9) and inhibitory neuron clusters (720 in Inh6), while upregulation in AD was observed primarily in oligodendrocyte clusters (949 in Oligo1). The largest number of DEGs upregulated in AD was observed in the Oligo7 cluster. However, this subtype represents a hybrid cluster, comprised of similar numbers of nuclei annotated as oligodendrocytes and excitatory neurons (42.4% and 38.4% of cluster nuclei, respectively). Thus, oligodendrocytes (Oligo7_Oligo) and excitatory neurons (Oligo7_Exc) in this cluster were analyzed separately for differential gene expression. Similar numbers of DEGs were identified for each of these subsets (2,530 for Oligo7_Oligo and 2,905 for Oligo7_Exc).

Biological pathway analysis of the PD-upregulated DEGs for each cell subtype showed the strongest enrichment in the Astro2 subtype, followed by other astrocyte, excitatory neuron, and oligodendrocyte clusters (Fig. 6Eii). These were dominated by pathways associated with neuronal morphogenesis/organization and synaptic transmission. Accordingly, the most strongly upregulated AD- and PD-GWAS genes were also involved in cell morphogenesis and organization, including *B3GAT1* in Astro2, and *GJC1*, *EFNA2*, and *PLK5* in Exc9 (Fig. 6Ei). Genes upregulated in AD over PD showed the strongest enrichment for pathways in the Oligo7 cluster (both Oligo and Exc subsets) as well as several other oligodendrocyte clusters (Fig. 6Eiii). Across these cell types, the top enriched pathways were largely associated with autophagy, mitochondrial structure, membrane trafficking, and mRNA processing. However, in Oligo1 and 7, the most strongly AD-upregulated individual GWAS genes were mainly associated with different pathways, including numerous protein synthesis and maturation-associated DEGs (Fig. 6Ei). These included ribosomal genes *RPS11*, *RPS15* and *RPL13A*, and chaperone *PFDN2* in Oligo1, and genes associated with cell cycle regulation (*FLBXL15*, *RPRML*), proteolysis (*FLBXL15*, *PSMC5*), and mitochondrial oxidative metabolism (*SLC25A39*, *CYC1*) in Oligo7.

To summarize, comparison of gene expression in DLB to either AD or PD primarily revealed gene upregulation in DLB within relatively few excitatory neuron and oligodendrocyte cell subtypes, but comparison of AD to PD revealed more diverse patterns of differential gene expression, with upregulation in PD within astrocyte, excitatory neuron, and inhibitory neuron clusters, and upregulation in AD within numerous oligodendrocyte clusters.

## 4. DISCUSSION

The three major NDDs AD, PD and DLB, are defined as distinct disorders but have common comorbidities, shared clinical presentation and overlapping pathological characteristics. In this study, we aimed to identify shared and divergent gene expression patterns among these NDDs at a granular cell subtype resolution. We thus compared the transcriptomic landscapes of AD, DLB, and PD within specific cell subtype populations of the TC. We utilized snRNA-seq datasets obtained from each of the three NDDs to gain insight into various aspects of pathogenesis across the different NDDs including: (1) vulnerability of specific cell subtypes, (2) disease-driver cell subtypes based on enriched expression of GWAS genes, (3) changes in cell-to-cell communication, (4) shared and (5) differential gene expression patterns and biological pathways (Fig. 1B).

NDDs are characterized by progressive neuronal loss. While vulnerable neuronal populations have been described for individual NDDs^46–48^, no previous work has directly compared vulnerability of the same cell subtypes across NDDs. We therefore examined depletion of excitatory and inhibitory neuronal subtypes in each NDD, and found that AD and DLB share a common vulnerable TC inhibitory neuron subtype. This neuronal type was characterized in part by expression of the major interneuron marker *VIP* and lack of expression of *PVALB*, *SST*, and *HTR3A*. Previous work has demonstrated cortical *VIP*^+^ interneurons to be moderators of cortical disinhibitory circuits, inhibiting *PVALB*^+^ and *SST*^+^ interneurons and thereby preventing inhibition of pyramidal neurons, thus regulating motor integration and cortical plasticity^49^. Loss of this subtype in AD and DLB suggests its potential involvement in cognitive impairment associated with both NDDs. In PD, previous work has primarily focused on characterization of vulnerable neuronal populations within the substantia nigra (SN)^48^. However, in this work we identified a cluster of inhibitory neurons depleted within the TC that was distinct from depleted populations in AD and DLB, suggesting potential association of this cell type with PD-specific pathology.

To better understand brain cell types driving disease risk in each NDD, we took a unique approach by examining enrichment of GWAS-gene expression. Multiple neuronal subtypes were implicated as disease drivers in DLB and PD, but in AD we identified only a single oligodendrocyte subtype. While published work has focused mainly on the role of disease-associated microglia in AD pathogenesis^50–52^, more recently the involvement of oligodendrocytes has also been suggested ^53,54^. Demyelination has been shown to often precede neuronal loss in AD cases^55^, and to result in neurodegeneration through disruption of metabolic axon support and maintenance^56^. Oligodendrocyte dysfunction causing myelin loss may thus represent a primary feature of AD pathology^57^. Furthermore, the importance of AD risk gene expression in oligodendrocytes has also been established^58^. For example, the major AD risk-associated gene *BIN1*, involved in vesicle endocytosis and regulation of apoptosis, among other functions, is primarily expressed in oligodendrocytes and has been implicated in AD-associated demyelination^59^. Here, we identified strong inhibition of *BIN1* in the disease-driver oligodendrocyte cluster of AD nuclei compared to other oligodendrocyte subtypes, along with highly increased expression of numerous other AD-GWAS genes associated with vesicle trafficking and apoptosis, including *PICALM* and *SNX1*. Dysregulation of these processes within disease-driver oligodendrocytes may contribute to oligodendrocyte dysfunction and AD progression within the TC.

Analysis of altered cell-to-cell communication also highlighted oligodendrocyte subtypes in all three NDDs, in addition to several neuronal subtypes. While in AD the strength of many communication pathways was increased, overall decreased communication was observed in DLB and PD. Together with our identification of the disease-driver cell types, these changes in cellular communication suggest an increased involvement of oligodendrocyte-neuron interaction in AD, while communication between and within these cell types may be inhibited in the context of the synucleopathies.

Here we also studied shared dysregulation of gene expression and impaired biological mechanisms across NDDs. We identified the highest numbers of shared DEGs among inhibitory neuron subtypes, most of which were downregulated in the NDD state. Previous studies have established an important role for inhibitory neurons in AD^60–62^, demonstrating that GABAergic neurotransmission is impaired both in human patients^63–65^ and murine AD models^66–68^, leading to hyperexcitability of neural circuits and likely contributing to cognitive dysfunction. In PD, it has been suggested that dysregulation of GABAergic neurotransmission is a primary driver of motor control deterioration^69^. Overaccumulation of intracellular Ca^2+^ along with *SNCA* is directly associated with neuronal death in PD in part through mitochondrial stress-induced apoptosis^70,71^, while GABA signaling prevents Ca^2+^ influx and thereby protects neurons from calcium toxicity^70^. Loss of dopaminergic neurons in the SN is furthermore predicted to dysregulate GABAergic neurotransmission^72,73^. These findings support the importance of inhibitory neurons in both cognitive decline in AD and motor deterioration in PD, as well as presumably in the combination of these clinical symptoms in DLB. Furthermore, our pathway analysis in inhibitory neuron subtypes revealed altered expression of numerous genes involved in mitochondrial processes across the NDDs, possibly indicating dysregulated metabolic activity resulting from disease-associated neurological dysfunction.

While NDDs share several molecular features and underlying mechanisms, each disease also displays unique molecular underpinnings associated with distinct biological pathways. We investigated the diseases-specific molecular determinants by direct comparison of differential gene expression between pathologies. This analysis produced several key discoveries. First, a relatively small number of cell subtypes displayed strong differential gene expression in DLB compared to either AD or PD. Moreover, in both these comparisons, almost all DEGs were upregulated in DLB and only few were upregulated in the either AD or PD. In contrast, when comparing AD vs PD, the majority of cell subtypes exhibited relatively high numbers of DEGs, with greater diversity in the directionality of differential expression across cell types. These observations indicate overall greater transcriptomic divergence between AD and PD than between DLB and either of the other NDDs, and support a model wherein DLB is positioned between AD and PD on a spectrum of neurodegenerative pathology.

In comparisons between all NDDs, we found that DEGs were predominantly identified in excitatory neuron and oligodendrocyte subtypes. Comparisons of PD to both AD and DLB identified multiple oligodendrocyte clusters with altered transcriptional profiles. Consistently, previous single-cell sequencing studies have revealed enriched expression of PD-GWAS genes in oligodendrocytes of the SN^74^, as well as depletion of differentiating oligodendrocytes in the midbrain of PD patients^75^. Furthermore, PD-specific oligodendrocyte populations have been predicted to display aberrant myelination activity based on human transcriptomic and mouse model data^76^. Together with these previous findings, our data suggest an important role for oligodendrocyte subtypes in PD that is distinct from both AD and DLB.

This work provides an essential direct comparison of the molecular underpinnings of three major NDDs. However, there are some limitations. First, in order to directly compare the transcriptomic signatures of the three NDDs, it was necessary to examine the same brain region in each context. However, brain regions are affected differently in each NDD. While neurodegeneration in cortical tissue may be associated with all three diseases, it is a hallmark only of AD and DLB, wherein dementia is an essential diagnostic feature. In PD, the TC region is typically involved in later stages of disease progression, when cognitive decline may occur^77–82^. In this work, the majority of PD donor samples were in earlier disease stages and exhibited little to no Lewy pathology within the TC, based on established metrics^83^. Thus, our data for PD reflect transcriptional changes preliminary to major neurodegeneration. Secondly, the relationships described here between gene expression and pathogenic mechanisms are predictive in nature and empirical validation through controlled experimentation in model systems is necessary to confirm the importance of these predicted mechanisms in the three NDDs.

Here we examined similarities and differences between the transcriptomic landscapes of three major NDDs. However, it is important to note that each of these disease categories represents a complex range of comorbid clinical symptoms and co-pathologies. Four major subtypes of AD have been characterized based on tau distribution, neurodegenerative patterns, and other pathological factors^84^. In addition, a recent multicentric study identified five molecular subtypes of AD using mass spectrometry proteomics of cerebrospinal fluids. Subtypes also differed in specific AD genetic risk variants, clinical outcomes, survival times, and patterns of brain atrophy^85^. Likewise, PD has been divided into three distinct subtypes based on both motor and non-motor factors including cognitive impairment, sleep disorder, and autonomic dysfunction^86^. DLB is particularly complex to define due to its shared clinical features with both AD and PD, but specific subtypes of this disease have also been described based on patterns of α-synuclein and tau distribution^87^. Future studies may thus apply similar strategies as are described here to elucidate the transcriptomic mechanisms underlying these pathological subtypes in order to develop an even higher-resolution understanding of the specific genetic factors driving diverse clinical outcomes. Because of the heterogeneity within and across NDDs, there is no single “silver bullet” for fighting neurodegeneration, but our findings provide unique predictive insight into the shared and distinct molecular mechanisms underlying these three pathologies, and contribute to a framework for future studies aimed at the development of targeted treatment strategies tailored to address the specific clinical challenges presented by each of these important diseases.

## ACKNOWLEDGEMENTS

We thank the Kathleen Price Bryan Brain Bank at Duke University (funded by NIH/NIA R01 AG028377) for providing us with the brain tissues, and the Duke Sequencing and Genomic Technologies Shared Resource for sequencing. This study used a high-performance computing facility partially supported by grant 2016-IDG-1013 (“HARDAC+: Reproducible HPC for Next-generation Genomics”) from the North Carolina Biotechnology Center.

## ETHICS APPROVAL AND CONSENT TO PARTICIPATE

The project was approved by the Duke Institutional Review Board (IRB). The study does not involve living human subjects. All samples were obtained from autopsies, and all are de-identified.

---

**Figure S1.**
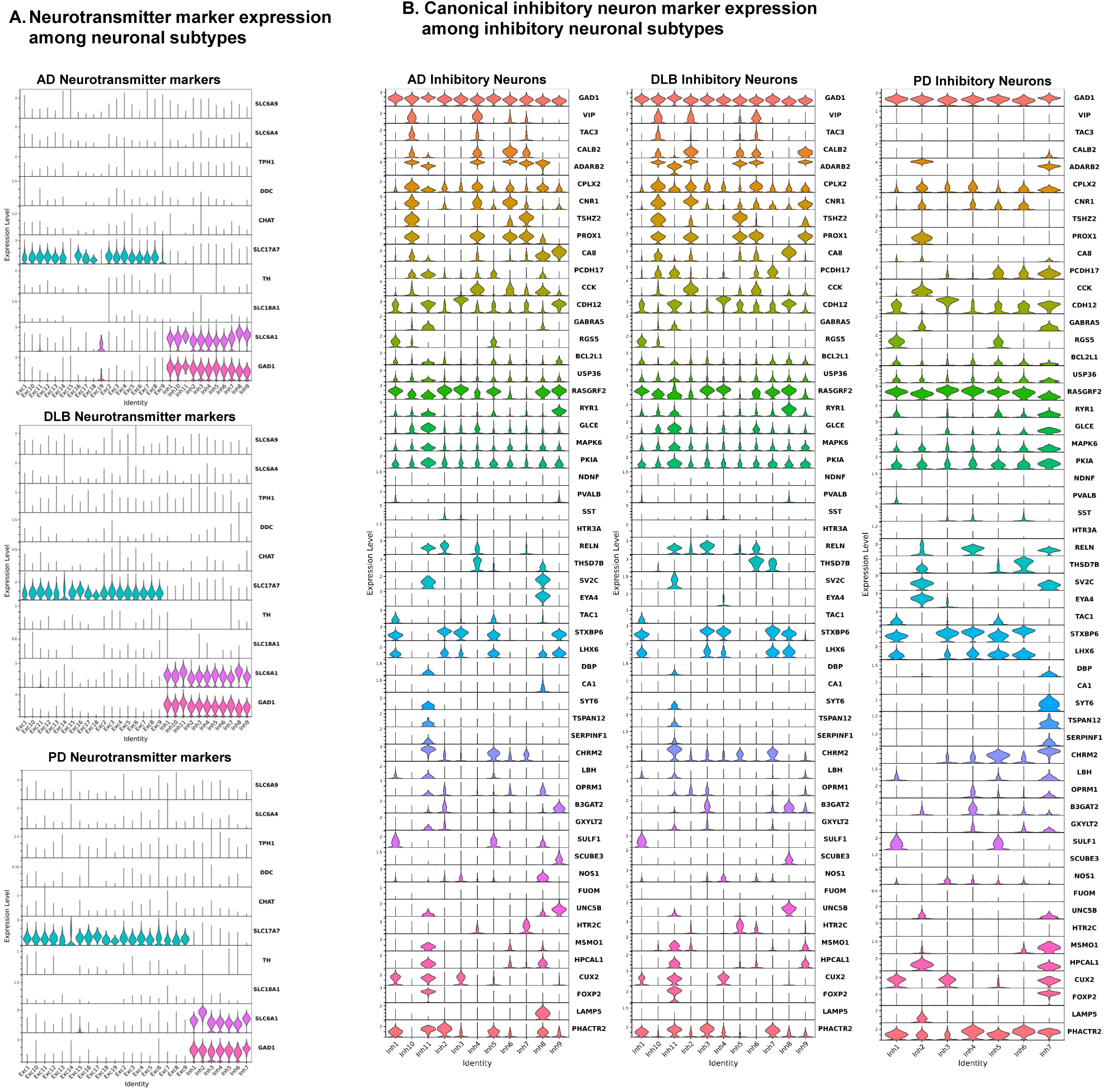
Canonical marker expression in neuronal subtypes of each NDD. A. Violin plots of log-normalized count data showing expression of canonical neurotransmission-type marker genes across neuronal subtype clusters of the three NDDs. Expression of 10 marker genes for neuronal subtypes engaged in signaling via different neurotransmitter molecules was examined. Inhibitory neuron clusters expressed genes indicating GABA transmission (*SLC6A1*, *GAD1*), while excitatory neuron clusters all expressed *SLC17A7*, indicating glutamate transmission. Other neurotransmission markers were not expressed in any of the clusters within the dataset, including markers for glycine transmission (*SLC6A9*), serotonin transmission (*SLC6A4*, *TPH1*), dopamine transmission (*DDC*, *TH*), acetylcholine transmission (CHAT), and general amine transmission (*SLC18A1*). B. Violin plots of log-normalized count data showing expression of expanded inhibitory neuron markers among all inhibitory neuron subtype clusters for each NDD.

**Figure S2.**
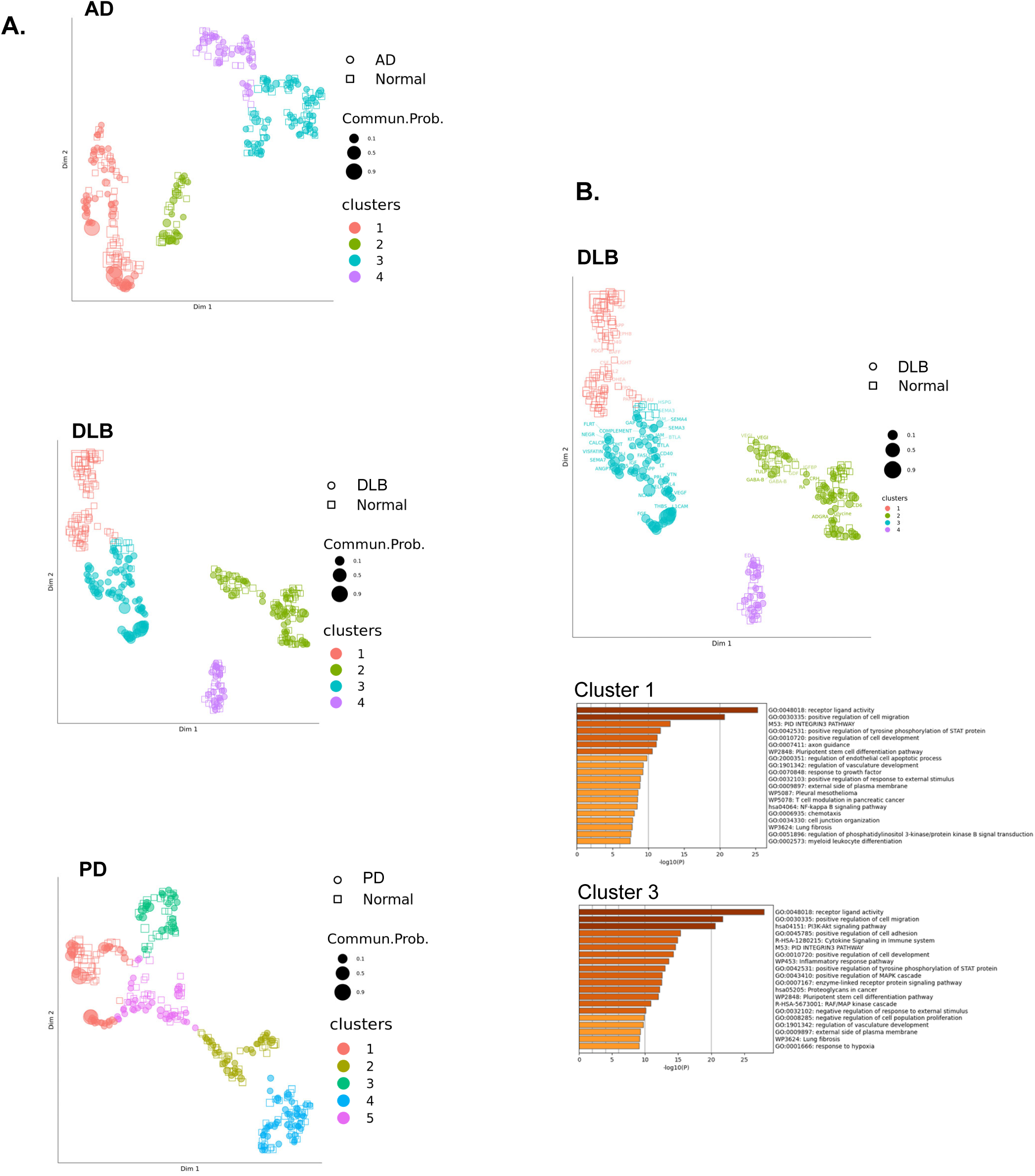
Clustering of communication pathways by interacting celltypes involved. A. Dimensional reduction and clustering of communication pathways based on transmitting and receiving cell types. Clusters of pathways based on similarity of interacting cell subtypes are color coded and numbered. Communication pathways in NDD nuclei are represented by colored circles and pathways of NC nuclei are represented by open squares. Point sizes are proportional to probability of communication. B. Metascape pathway analysis of top 20 enriched biological pathways among genes involved in interactions between celltypes in DLB communication clusters 1 (Normal dominant) and 3 (DLB dominant).

